# Evolutionary sequence and structural basis for the distinct conformational landscapes of Tyr and Ser/Thr kinases

**DOI:** 10.1101/2024.03.08.584161

**Authors:** Joan Gizzio, Abhishek Thakur, Allan Haldane, Carol Beth Post, Ronald M. Levy

**Affiliations:** Center for Biophysics and Computational Biology, Temple University, Philadelphia, Pennsylvania 19122; Department of Chemistry, Temple University, Philadelphia, Pennsylvania 19122; Department of Physics, Temple University, Philadelphia, Pennsylvania 19122; Borch Department of Medicinal Chemistry and Molecular Pharmacology, Purdue University, West Lafayette, Indiana 47907

## Abstract

Protein kinases are molecular machines with rich sequence variation that distinguishes the two main evolutionary branches – tyrosine kinases (TKs) from serine/threonine kinases (STKs). Using a sequence co-variation Potts statistical energy model we previously concluded that TK catalytic domains are more likely than STKs to adopt an inactive conformation with the activation loop in an autoinhibitory “folded” conformation, due to intrinsic sequence effects. Here we investigated the structural basis for this phenomenon by integrating the sequence-based model with structure-based molecular dynamics (MD) to determine the effects of mutations on the free energy difference between active and inactive conformations, using a novel thermodynamic cycle involving many (n=108) protein-mutation free energy perturbation (FEP) simulations in the active and inactive conformations. The sequence and structure-based results are consistent and support the hypothesis that the inactive conformation “DFG-out Activation Loop Folded”, is a functional regulatory state that has been stabilized in TKs relative to STKs over the course of their evolution via the accumulation of residue substitutions in the activation loop and catalytic loop that facilitate distinct substrate binding modes *in trans* and additional modes of regulation *in cis* for TKs.

## Introduction

Serine/threonine protein kinases (STKs)^1^ are very ancient and ubiquitous signaling enzymes; despite their common name “eukaryotic protein kinases”, these protein domains are also present in archaea and bacteria ^2^ suggesting their presence in the last universal common ancestor (LUCA) 3-4 billion years ago.^3^ Eukaryotic tyrosine kinases (TKs) appear to have descended from this lineage much later on, just prior to the emergence of the first metazoans ∼600 million years ago ^4–6^. TKs share a great deal of structural homology with even the most distantly related STKs. Notable differences however can be seen in catalytic domain motifs that specialize TKs for phosphorylating tyrosine instead of serine and threonine, in addition to unique domain organizations that make them critical for multicellular function; the majority of TKs are fused to a receptor domain which directly facilitates signaling between cells.^7^ TKs comprise less than 20% of the ∼498 kinases in the human genome^8^, yet these proteins are among the most frequently mutated genes in cancers ^9^ emphasizing their important role in multicellular function. Consistent with this view, TKs with catalytic domains homologous to those in humans are also extensively found in ancient holozoan lineages that display animal-like multicellular behavior such as choanoflagellates.^6,10^

The catalytic domains of both TKs and STKs contain mobile structural elements that allow the active state of the enzyme to be dynamically assembled and disassembled, at the cost of a free-energy penalty that depends on their amino acid sequence; the activation loop, glycine-rich loop (also called G-loop or P-loop), and αC-helix each contain at-least one conserved residue that is directly involved in the catalytic process for all kinases in the “active conformation”. Kinase catalytic domains display a large degree of sequence variation at positions which, when viewed in the tertiary structure, are located at intra-molecular interfaces involving these mobile structural elements, especially the activation loop where the sequences of TKs and STKs are highly diverged.^11,12^ The activation loop is a flexible ∼20 residue-long motif beginning at the conserved DFG (Asp-Phe-Gly) residues, where the DFG-Asp plays a direct role in catalysis. The primary sequence of the loop terminates just before the conserved APE (Ala-Pro-Glu) residues in the enzyme’s C-terminal lobe; this C-terminal region of the activation loop (also referred to as P+1 loop in the literature) forms part of the conserved inter-molecular binding surface for Ser, Thr, or Tyr residues of peptides and proteins targeted for phosphorylation, although it is shaped very differently in TKs compared with STKs which reflects their evolved specificity for Tyr vs Ser/Thr substrates ^11,13^. In the active conformation, the activation loop interacts non-covalently with the catalytic loop region at two ends via its N- and C-terminal “anchors”^13^. The catalytic loop, like the activation loop, also contains catalytic residues, e.g., the His-Arg-Asp (HRD) motif, but the catalytic loop is comparatively rigid in structure allowing it to serve as a central hub for the organization of the active conformation. There is yet a third anchor point for the activation loop, the “RD-pocket” ^13^, which is centered on the HRD-Arg sidechain and packs against residues of the activation loop N- and C-terminus. The RD-pocket is a positively charged surface formed by a cluster of basic sidechains which are stabilized by the covalent addition of acidic phosphates to activation loop Ser, Thr, or Tyr residues ^14^. The sequences of TKs and STKs differ significantly at all three anchors, which was suggested by our previous computational and structural analysis to result in very different conformational free energy landscapes on-average between TKs and STKs.^15^ These anchor points also interact directly with substrate peptides which bind in distinct ways to alternative surfaces surrounding the shared active site of TKs and STKs^16^, and therefore the evolutionary divergence in substrate specificity may be intrinsically coupled with that of the conformational landscape.

In living cells, the population of the active conformation relative to the inactive conformations is regulated by the intrinsic free-energy landscape of the catalytic domain as well as external factors such as post-translational modifications, protein-protein binding, domain assemblies, etc.^17^ These inactive conformations can be viewed as “basins” of the free-energy landscape in a very high dimensional conformational space. Characterizing these basins computationally or experimentally is challenging, but over the decades there have been thousands of kinase structures in the active and inactive conformations characterized with x-ray crystallography and deposited in the Protein Data Bank (PDB), allowing relevant structural degrees of freedom to be mapped.^18–22^ A variety of inactive conformations have been observed in x-ray crystallographic and solution NMR structures involving various disruptions of the catalytic machinery, some more drastic than others in the extent of activation loop displacement. Among these examples are structures of inactive kinases displaying a “classical DFG-out” conformation^18^ where the activation loop backbone is reorganized by ∼20 Å relative to the active-extended state with the N-terminal anchor completely dislocated. This large backbone reorganization appears to propagate from a “flip” of the DFG motif at the beginning of the activation loop from “DFG-in” to “DFG-out”, and in this way downstream residues are placed in proximity to a peptide binding surface in the C-lobe which has been functionalized by TKs.^13,16,17^ Previously it was confirmed ^15^ that the stability of this “**DFG-out activation loop-folded**” basin for TKs relative to STKs is greatly enhanced by contacts *in cis* that allow the folded/inactive activation loop to mimic a Tyr-containing protein substrate that would otherwise bind to the active conformation *in trans*^17,23^. This appears to be a complex form of TK autoregulation in which the very selection pressures that have optimized TKs for binding tyrosine substrates also stabilize the substrate-competitive DFG-out activation loop-folded conformation. The substrate-mimicry of the activation loop is a notable feature of this basin that sets it apart from alternative, minor activation loop reorganizations; for example, the Src-like inactive conformation which involves a large rotation of the αC-helix but only “partial” folding of the activation loop and, for TKs, appears to be an intermediate state along the transition pathway to DFG-out^22,24^. Some inactive conformations of TKs observed in the PDB display a flip of the DFG motif from “in” to “out”, but the rest of the activation loop is minimally perturbed relative the active state, displaying an extended activation loop that is readily capable of binding substrates (e.g., PDB: 2G2F)^22^. This DFG-out conformation where the activation loop remains extended is a tractable target for biased and unbiased MD simulations of active ↔ inactive conformational transitions for TKs.^25,26^ However, recent experimental^27^ and computational evidence^15^ suggests the DFG-out conformation with a fully folded activation loop is the major DFG-out inactive state of TKs in the absence of inhibitors, and is further stabilized by the binding of type-II inhibitors^27,28^. Despite this, there remains a tendency in the computational literature to focus on conformational changes of TKs involving a localized “DFG flip” between “in” and “out” without the large-scale reorganization of the activation loop between “extended” and “folded”^25,29^.

Previously we estimated the reorganization free-energy (Δ*G*_*reorg*_) for apo STK catalytic domains to undergo the large-scale active DFG-in → inactive DFG-out conformational change of the activation loop from extended to folded, which we estimated to be 4-6 kcal/mol larger on average than for TKs.^15^ This estimate came from a combination of sequence-based “threading” with a machine-learning statistical energy function and a structure-based indirect method using the binding free energies of type-II inhibitors estimated from FEP simulations involving the inactive DFG-out conformation as a computational probe to extract the reorganization free-energy contribution from experimental binding affinities reported in the literature. The sequence-based approach which we have employed in both the previous and current work is based on a maximum-entropy Potts statistical energy model that parameterizes the fitness of aligned kinase sequences in terms of direct residue-residue interactions.^30^ Provided an MSA of sufficient depth, the statistical energy parameters for all possible combinations of residue pairs can be determined using a generative MCMC procedure implemented by the Mi3-GPU software.^31^ Through repeated MCMC runs the Mi3-GPU inference algorithm iteratively converges on a network of couplings that are capable of generating sequence ensembles with residue correlations matching what is observed in the protein family MSA. The relationship between Potts statistical energies and fold stability has long been recognized^32,33^; this has motivated the use of Potts models in the study of protein folding landscapes^34^ and the effects of mutations on fold stability^35,36^ but the extent to which the relative likelihood of occupying individual conformational basins that comprise the folded landscape are captured by the Potts model is a newly developing area of study. Our previous estimates of the relative fold stability of TKs vs STKs in the active vs inactive conformations were obtained by threading the Potts coupling terms onto active and inactive structures sampled from the two conformational basins.^15^ This provided a decomposition of the energy landscape in terms of residue-residue interactions which enabled us to identify structural motifs that are responsible for the observed divergence in Δ*G*_*reorg*_between TKs and STKs for the conformational change where the activation loop reorganizes from the extended (active) state to the folded (inactive) and forms distinct substrate binding surfaces occupied by peptides when the kinase is active.

In this current study we sought a direct verification of the correspondence between Potts threaded energies and reorganization free energies of TKs and STKs and the structural motifs which, in nature, have diverged in sequence to “tilt” their active → inactive landscapes in opposite directions. Using mutation FEP simulations and a novel thermodynamic cycle involving two different native protein environments, the effects of activation loop anchor point mutations on the relative stability of the active (DFG-in, activation loop extended) and inactive (DFG-out, activation loop folded) states were calculated. We compared the effects of 54 mutations predicted by the Potts model to result in the largest change in relative stability of the two basins (active vs. inactive) by performing 108 FEP simulations to alchemically mutate sidechain(s) in both the active DFG-in and autoinhibited/”activation loop folded” DFG-out conformations. As described below, our results confirm the correspondence between the Potts statistical energy changes in sequence space and conformational free-energy changes in structure space and support previous hypotheses bout the structural and sequence basis for the evolutionary divergence in conformational landscapes of TKs compared with STKs. By using the Potts model and FEP simulations to directly mutate the activation loop anchor residues of STKs to residues commonly seen in TKs (and vice versa), the role of sidechain interactions in the active basin that form the distinct substrate binding surfaces of TKs and STKs were confirmed to destabilize the active DFG-in basin in TKs relative to STKs. Regarding the inactive basin, the Potts model and FEP simulations recapitulate previous insights into the importance of pseudo-substrate interactions between the activation loop and its own substrate binding surfaces for stabilizing the inactive conformation in TKs.^17,23,37,38^

## Results

### 1. Conformational preferences predicted by FEP match Potts sequence-based predictions

Potts statistical energy models can be used to compare the relative stability of individual conformational basins that comprise the native ensemble for individual sequences.^15,39^ The sequence-based statistical energy predictions of relative conformational stability can then be directly compared with thermodynamic observables calculated from all-atom MD free energy simulations. Here we describe and validate the consistency of these two distinct approaches, which we leverage to investigate the sequence and structural basis for the evolutionary divergence in an “active → inactive” conformational change for TKs versus STKs.

#### 1.1. Choosing kinases for targeted analysis of conformational differences between TKs and STKs

The following analysis required experimental structures of kinases which have both an active structure in the PDB where the activation loop is “extended”, and an inactive structure in a classical DFG-out conformation with an autoinhibitory “folded” activation loop (Figure 1). For conciseness, we refer to these conformational states as “active” and “inactive”, respectively.

**Figure 1.**
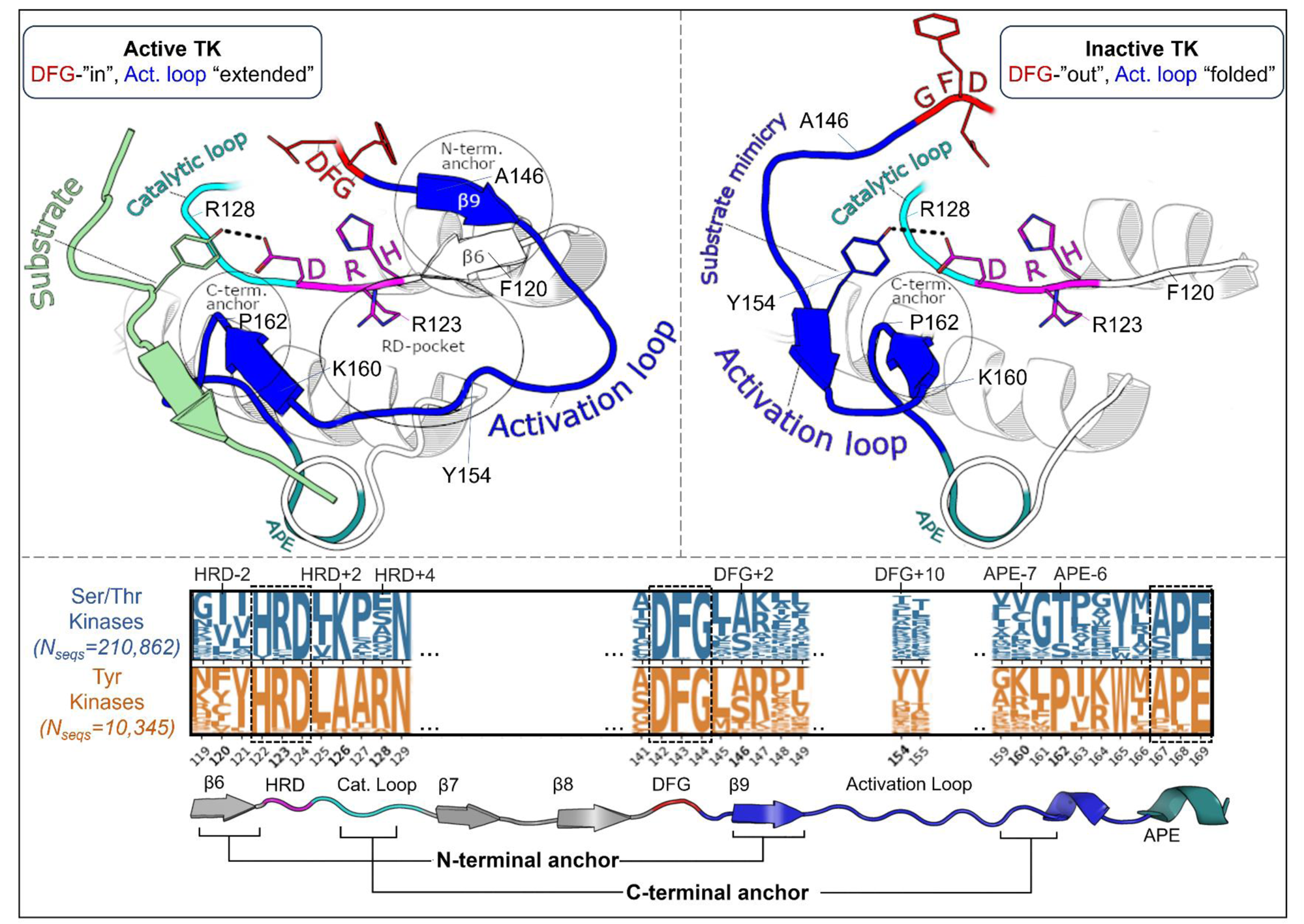
Overview of the kinase active and inactive conformations and sequence features that distinguish TKs (Tyr Kinases) from STKs (Ser/Thr Kinases). (top) Illustrations of the active and inactive conformation of the activation loop for a representative TK, INSR. The active conformation is characterized by three main structural motifs – the N-terminal anchor, RD-pocket, and C-terminal anchor. In the inactive conformation, the C-terminal anchor of TKs remains intact while the rest of the activation loop is “folded up”, with the DFG+10 residue Y154 mimicking the binding mode of Tyr substrates. (bottom) Sequence logo visualization of key motifs in our MSA of kinase catalytic domains, where the vertical axes represent the raw residue frequency for STKs (blue) and TKs (orange) in our MSA, calculated separately. The residue numbering in this MSA is displayed on the horizontal axis. Only key motifs are displayed for clarity. The conserved triads “HRD”, “DFG”, and “APE” were used as reference points for a more general numbering scheme: for example, DFG+10 refers to residue 154 in our alignment (written as 154_DFG+10_ in the main text) and is located ten residues C-terminal from the DFG motif, and the Gly of DFG is located at position 144 in our MSA. Sequence logos were plotted using Logomaker^40^.

To analyze the evolutionary divergence in the active → inactive free energy landscapes of TKs versus STKs, three kinases were chosen with structures in both conformations: INSR which is a TK, and two STKs, BRAF and CDK6 which belong to the TKL and CDK families, respectively (Figure 2, top). The active, extended conformation (Figure 1, top left) of the activation loop is characterized by three anchor points:

i. “N-terminal anchor” ^13^ – a structural motif formed by a pair of antiparallel β-strands at the activation loop N-terminus (β9: residues 145_DFG+1_ through 147_DFG+3_) and the N-terminal region of the catalytic loop (β6: residues 119_HRD-3_ through 121_HRD_). TKs and STKs have notable sequence differences in the β6 strand.
ii. “RD-pocket” ^13,14^ – a cluster of basic sidechains involving the activation loop (residues 147_DFG+3_, 154_DFG+10_ and/or 155_DFG+11_) and Arg of the conserved HRD motif (residue 123_HRD_). Residue 160 from the activation loop C-terminal region is tightly packed against R123_HRD_ in STKs but decoupled from the RD-pocket in TKs.
iii. “C-terminal anchor” ^13^ – a structural motif characterized by non-covalent interactions between the C-terminus of the activation loop and the catalytic loop. The C-terminal anchor forms a binding surface for Ser, Thr, or Tyr substrates.

**Figure 2.**
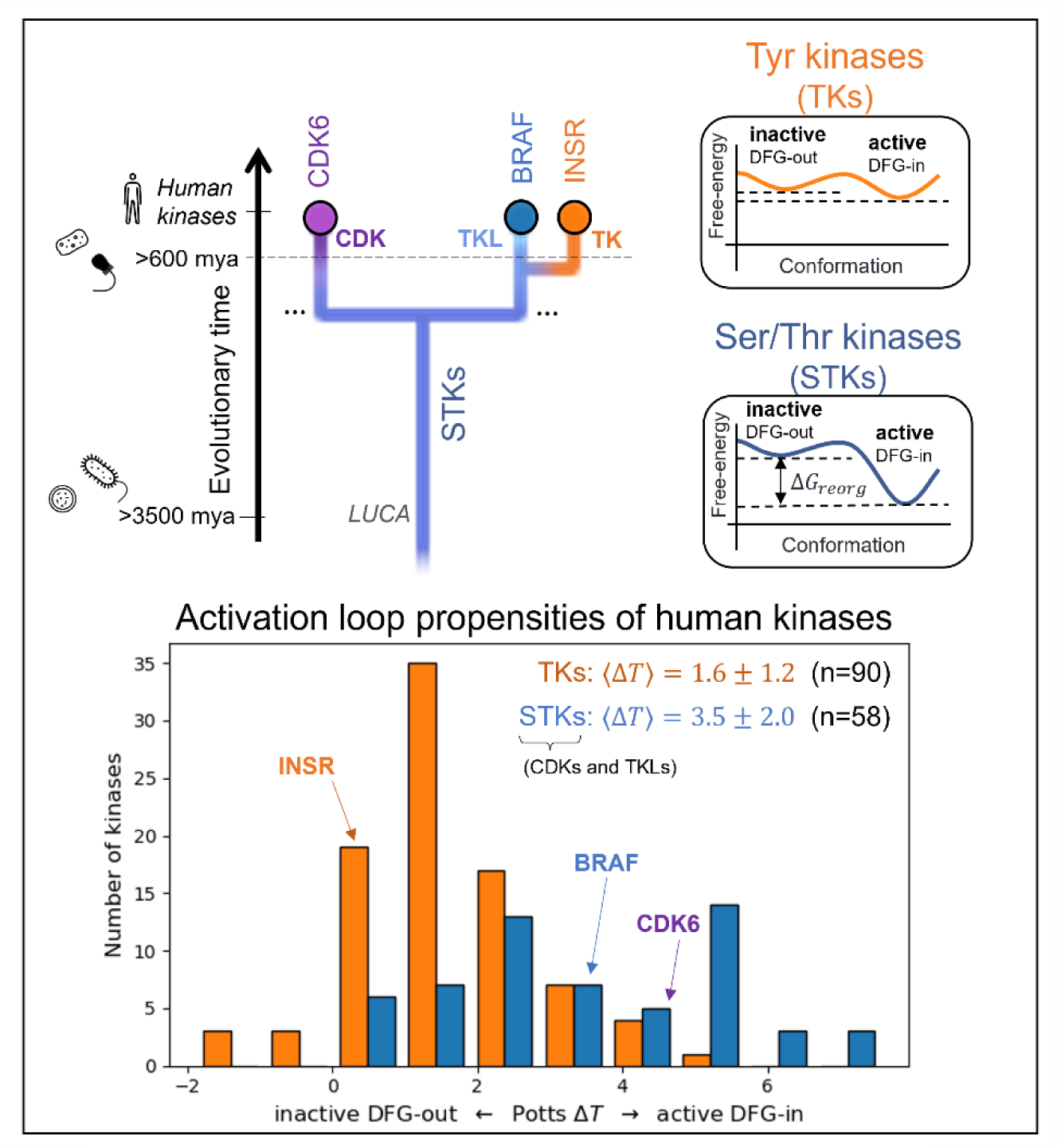
Choosing kinase targets (top) to analyze differences in the conformational landscapes of TKs and STKs. CDKs are STKs belonging to the “CMGC” group^8^, which are distantly related to TKs. BRAF, an STK from the “Tyrosine Kinase-Like” (TKL) group, is more closely related to TKs. INSR is a typical receptor TK, with active and inactive conformations that are representative of both cytoplasmic and receptor TKs. Icons are displayed to the left to mark the divergence of kinase-containing taxa (from top to bottom – H. sapiens^8^ and porifera^41^ (animals), choanoflagellates^10^, archaea and bacteria^3^). The Potentials of Mean Force (PMFs) shown on the right are artistic illustrations to help visualize the free-energy landscapes of TKs and TKs and STKs described previously^15^ – while the barrier heights are unknown, the relative depths of the two basins at the end-states are accurately depicted. (*bottom*) The Potts DFG-out penalties of 90 human TKs (receptor and non-receptor families) and 58 STKs (41 TKL kinases and 21 CDKs) were estimated by threading over structural ensembles of the active and inactive (DFG-out) conformations (see *Methods*), showing a bias for human TKs (orange) towards inactive relative to STKs (blue).

All three of these motifs (*i-iii*) may be disassembled in the inactive conformation where the activation loop is folded. However, TKs are unique in that they maintain an intact C-terminal anchor in the inactive conformation. This allows a Tyr residue from the middle-segment of the activation loop (e.g., Y154_DFG+10_) to occupy the substrate binding site *in cis*^17^, a phenomenon we refer to as “substrate mimicry” (Figure 1, *right*).

#### 1.2. Potts threaded-energy prediction of conformational preference

Next we establish, based on Potts calculations, that TKs have an increased bias for the inactive state relative to STKs. For an aligned protein sequence (*S*), a corresponding protein family Potts model (in this case, the kinase superfamily) can be used to compare the stability of two conformational ensembles, e.g., active (labeled *A*) and inactive (labeled *B*), by “threading” that sequence’s statistical energy couplings 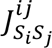 over pairs of residues *i* and *j* which change their contact frequency *c*^*ij*^ between the two ensembles,

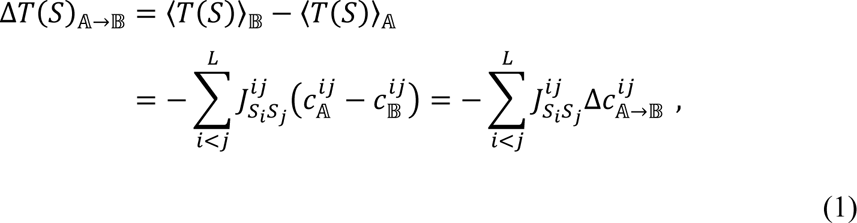

where *S* denotes a single kinase domain sequence aligned to a MSA of length *L* (number of columns). Δ*T* can be considered a Potts statistical energy analog of the reorganization free-energy Δ*G*_*reorg*_, a thermodynamic quantity that reflects the relative free-energy of the two conformational basins,

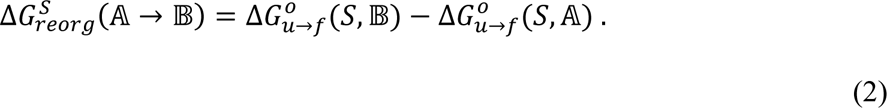

In this expression 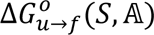 is the free energy change associated with the “protein unfolded → protein folded” transition where the definition of “folded protein” is restricted to a particular conformational basin *A* (**active DFG-in, activation loop extended**), and likewise for 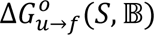 which instead restricts the folded state definition to basin *B* (**inactive DFG-out, activation loop folded**).

MD simulations of the three target kinases in their active and inactive states, CDK6, BRAF, and INSR, were used to calculate contact frequencies (see *Methods*), and these contact frequencies were used to thread homologous kinase sequences from the three families (human TKs, TKLs, and CDKs) using Equation (1). Plotted in Figure 2, the Δ*T* distributions for these families recapitulate our previous observations over a much larger set of sequences^15^ that, on-average, TKs are intrinsically more balanced between the active → inactive conformational basins due to common features of their sequences that distinguish them from STKs (including TKLs and CDKs, among other groups). Specifically, we found that the average difference in Potts threaded energy scores between TKs and STKs is approximately 4 kcal/mol when scaled to physical free-energy units.^15^ At this scale, the most populated bin in the Δ*T* histogram of TKs from Figure 2 (Δ*T* ≈ 1) is approximately 1.3 kcal/mol. This corresponds to a 1:10 ratio of inactive:active conformers in solution (10% inactive), consistent with experimental NMR populations for an isolated TK catalytic domain^27^ (7% inactive).

#### 1.3. Potts-generated library of mutational effects on conformational bias

Proteins can explore functional space by accruing mutations that shift the conformational equilibrium of the folded protein between different native-like conformations.^12,42,43^ This can be caused by changes in non-covalent interactions involving the mutated residue and nearby sidechains in the two different protein environments (e.g., active vs inactive conformations) and changes in the physicochemical nature of these interactions upon mutation. In terms of Potts threaded energy, we define the net mutational effect on the conformational preference as ΔΔ*T*,

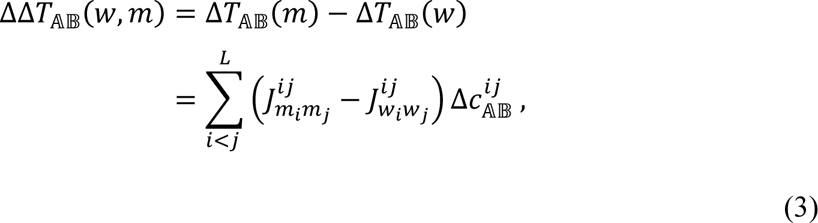

where *w* and *m* denote two distinct sequences of length *L* which differ at one or more positions. Only positions *i* or *j* which are “mutated” in sequence *m* relative to sequence *w* contribute to ΔΔ*T* (the net shift in Δ*T*), depending on how the threaded couplings of these positions with other residues in the protein are changed upon mutation. In this way, the Potts model can be used to scan for mutations which have significant effects on the conformational free-energy landscape.

For our target TKs and STKs (CDK6, BRAF and INSR) we scanned for mutations in the Potts model that are predicted by Equation 3 to induce the largest changes in conformational preference for the active vs inactive state (Figure 3A) and find that the largest projected mutational effects are mostly observed in association with structural motifs that are important for distinguishing the functional landscapes of TKs vs STKs.^15^ To predict mutations with the most significant effects on the active (*A*) ⇌ inactive (*B*) conformational equilibrium of TKs and STKs, ΔΔ*T* (Equation 3) was evaluated for ≈ 10^4^ mutations of STKs and TKs, using CDK6, BRAF and INSR as representative systems.

**Figure 3.**
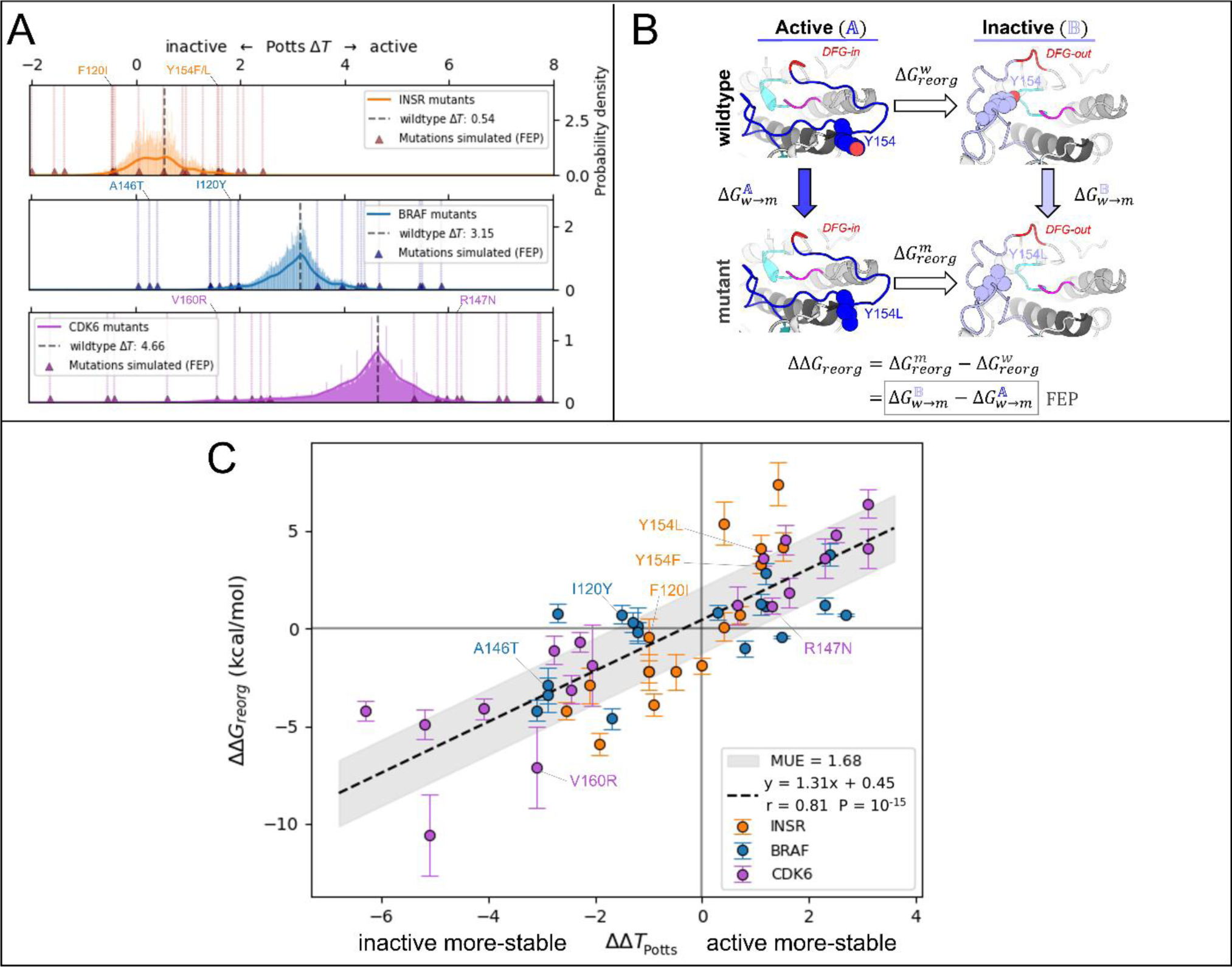
Potts statistical energy ΔΔ*T*s for the effects of mutations are consistent with corresponding ΔΔ*G*_*reorg*_from FEP. (**A**) Results of scanning double and single mutations in the Potts model, plotted as a histogram of raw mutant Δ*T*s (Δ*T*_*mut*_ = ΔΔ*T* + Δ*T*_*wt*_) for > 10^4^ mutations from each kinase (see *Methods*). Mutations chosen for FEP simulations are marked with triangles and scored with vertical lines. (see Table S3 for a full list of mutations chosen for FEP). (**B**) Thermodynamic cycle (left) which we used to calculate each of the 54 *ΔΔG*s in A. The vertical legs represent the alchemical transformations performed in FEP simulations in the active basin *A* and the inactive basin *B*, while the horizontal legs represent the physical free energy of reorganization between the two basins for wildtype (top) and mutant (bottom). (**C**) Plot of *ΔΔT* calculated from the Potts model vs *ΔΔG*_*reorg*_for 54 mutations calculated from 108 FEP simulations in the active and inactive conformations (see *Methods*). *ΔΔG*_*reorg*_and *ΔΔT* share a sign convention; positive values indicate a shift in conformational stability towards the active conformation, and negative values indicate a shift towards the inactive conformation.

#### 1.4. FEP predictions of mutational effects on conformational stability

To validate the Potts model predicted effects of these mutations, FEP simulations were employed using a novel thermodynamic cycle to calculate ΔΔ*G*_*reorg*_ (Figure 3B) – a structural thermodynamic observable for which ΔΔ*T* is a sequence-based analog.^14^ ΔΔ*G*_*reorg*_ represents the change in free-energy difference between active (*A*) and inactive (*B*) that occurs upon mutation, and can be calculated by alchemically mutating the target residue(s) in-place within the two conformational basins:

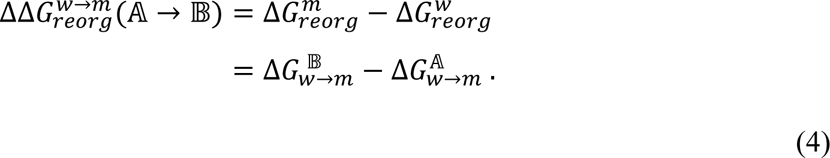

In this notation, the subscript of Δ*G* indicates the free-energy change associated with an equilibrium process, e.g., protein conformational reorganization or mutation of a sidechain via alchemical FEP simulations, and the superscript identifies the protein sequence (wildtype or mutant) or protein conformation (basin *A* or *B*) that the process is performed on. It is technically possible to use the first line of Equation (4) to calculate ΔΔ*G*_*reorg*_, but in practice the free-energy term Δ*G*_*reorg*_is very difficult (if not impossible) to converge from direct MD free-energy simulations of the activation loop reorganization along a physical path. When using the second line to calculate ΔΔ*G*_*reorg*_, only a structural perturbation of the sidechain in two different protein environments, basins *A* and *B*, is required (Figure 3B). Hence, the alchemical approach is more reliable and computationally efficient, provided the physical end states are stable at the simulated timescales or the appropriate restraint procedures are deployed.

#### 1.5. Correspondence of sequence- and structure-based methods

By calculating ΔΔ*G*_*reorg*_ for landscape-altering mutations and comparing with the sequence-based Potts model analogue ΔΔ*T* (Figure 3C), we were able to confirm that the structural basis for the divergent conformational landscapes of TKs and STKs originates from residues that are associated with their distinct substrate binding functionalities. As described in *Methods,* 56 mutations with the largest magnitude ΔΔ*T*s were selected for comparison with ΔΔ*G*_*reorg*_from FEP simulations, resulting in 108 free-energy simulations for mutations of TKs and STKs in the active and inactive conformations. As shown in Figure 3C, results from the sequence based / Potts and structure based / FEP approaches are highly consistent (Pearson correlation of 0.81 with p-value ≈10^−15^).

The slope of the linear fit can be considered an approximate conversion factor from statistical energy units to kcal/mol which from Figure 3C is ≈1.3 kcal/mol, close to our previous estimate^15^. This suggests the Potts coupling terms in Equation 3 are capturing physical information about the free-energy balance of sidechain interactions between two different conformational basins. This correspondence is noteworthy considering the Potts Hamiltonian is a strictly sequence-based information theoretic model, i.e., the potential function is not explicitly trained on structural information. The only structural information used in the Potts threading calculation is the definition of active and inactive macrostates from PDB or MD ensembles, and the determination of residue-residue contacts that break and form between the two ensembles (Δ*c*^*ij*^, see *Methods*). It is the threaded energy terms from Equation 3 that provide information about how the two distinct protein environments, active and inactive, respond differently to mutations.

### 2. Confirming the structural basis for the evolutionarily divergent conformational landscapes of TKs vs STKs

The mutations with the largest effects on the active → inactive free-energy landscape from Figure 3C tend to cluster in structural motifs in the active and inactive conformations that were previously identified to shape the distinctive landscapes of TKs compared with STKs due to sequence differences.^15^ Mutations of these motifs can increase the bias in the free-energy landscape towards the active conformation, characteristic of STKs, or decrease the relative stability of the active conformation, a signature phenotype of TKs.

In the active conformation, we find there are three main structural motifs which, due to average sequence differences between the two clades, broadly contribute to the divergent conformational landscapes of TKs vs STKs: the N-terminal anchor, RD-pocket, and C-terminal anchor. Here, we describe the structural basis for these effects on the stability of the active conformational basin, focusing on prominent examples of mutations which introduce a TK-prevalent residue into a wildtype STK sequence (CDK6), and test the effect of mutations at these locations using the Potts model and FEP. By comparison of observed residue positioning and physical interactions in PDB structures of TKs versus STKs, we establish structural rationales for the predicted mutational effects.

#### 2.1. The N-terminal anchor is destabilized in the active conformation in TKs relative to STKs

The N-terminal anchor (Figure 4A) is a motif formed in the active-state by a pair of antiparallel β-strands involving the activation loop N-terminus (β9) and the stretch of three residues located N-terminal to the HRD motif (β6).^13^ The N-terminal anchor can be identified by sequence positions 120 (HRD-2) and 121 (HRD-1) in the β6 strand, and 146 (DFG+2) through 148 (DFG+4) in the β9 strand in our MSA numbering (see Table S2 for equivalent PDB residue numbering). This strand pairing only occurs when the activation loop is “extended” as in the active conformation.^13,44^

**Figure 4.**
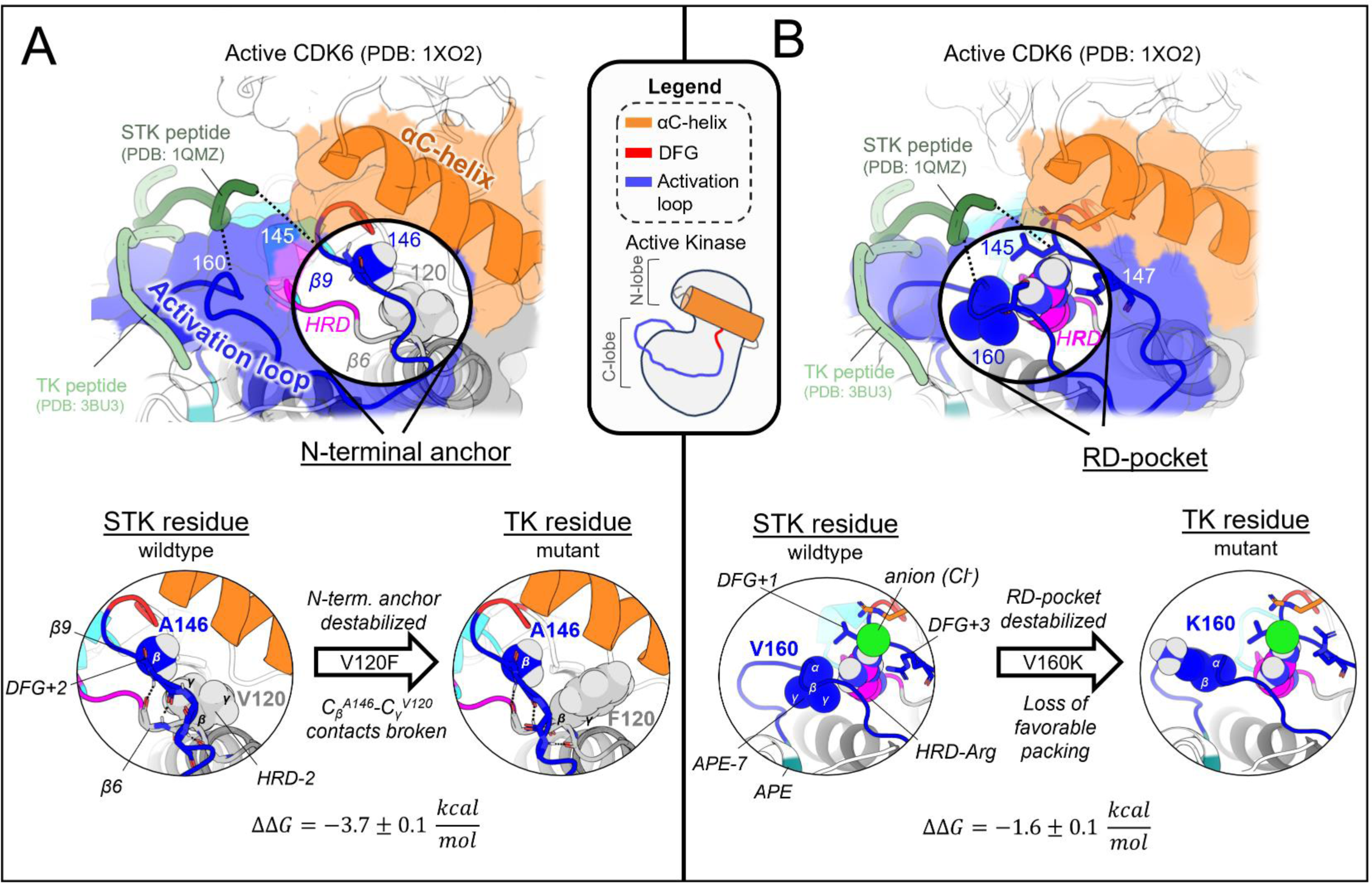
Mutating STK residues to those found in TKs destabilizes the active conformation relative to inactive. (**A**) *Top*: surface representation of wildtype CDK6 (PDB: 1XO2 chain B) in the active conformation (see Methods), with the N-terminal anchor highlighted. Peptides from TK (light green) and STK (dark green) holoenzymes are superimposed for reference – STK holoenzymes rely on interactions between peptides and the N-terminal anchor, in contrast to TKs which bind peptides further away. *Bottom*: van der Waals (vdW) space-filling models of the CDK6 wildtype (left) and mutant (right) N-terminal anchor residue 146_DFG+2_ and its interaction partner 120_HRD-2_, showing the loss of favorable vdW contacts between the C_β_ atom of 146_DFG+2_ and Cγ of 120_HRD-2_. Backbone hydrogen bonding patterns that define the β-strands are shown with dashed lines. (**B**) *Top*: same as *A* but with the RD-pocket highlighted. As before, peptides from TK and STK co-crystal structures are superimposed for reference. *Bottom*: vdW space-filling models of residue 160_APE-7_ in the wildtype (left) and mutant structure (right), located in the activation loop C-terminus, which stabilizes the RD-pocket in STKs. Small aliphatic residues like V160_APE-7_ in CDK6 pack favorably against the HRD-Arg, while bulky sidechains, e.g., K160_APE-7_ (seen in TKs), decouple from the RD-pocket and flip “out” in our MD free-energy simulations to interact with solvent (or peptides in the TK holoenzyme only).

In STKs, the N-terminal anchor is located inside the “cleft” between the N-lobe and C-lobe where residues of the holenzyme interact with bound peptides and protein substrates in co-crystal structures^16^ as illustrated in Figure 4A. By contrast, substrates co-crystallized with TK holoenzymes adopt a different orientation which places residues of the bound substrate further away from the N-terminal anchor.^16^ Consistent with this paradigm, our results based on MD free energy simulations and Potts calculations described next suggest the N-terminal anchor (β6-β9 pairing) evolved under weaker stability constraints in TKs compared with STKs. The β6 and β9 strands contain divergent sequence features which, as shown in TKs is relatively unstable compared with STKs.

Cross-strand sidechain interactions between position 120_HRD-2_ in the β6 strand and 146_DFG+2_ in the β9 strand are diverged between TKs and STKs due to sequence differences^15^; 77% of STKs in our MSA have, simultaneously, a branched aliphatic sidechain at 120_HRD-2_ (Val or Ile) and a small unbranched sidechain at 146_DFG+2_ (Ala or Ser), whilst fewer than half of TKs (39%) contain these residue pairs. Many TKs (23%) instead have a large aromatic sidechain (Phe) at β6/HRD-2, which by contrast is very rare for STKs (4%) and predicted by the Potts model to destabilize the N-terminal anchor. Specifically, when using the Potts model to mutate the β6-strand from V120_HRD-2_ to Phe in CDK6, or I120_HRD-2_ to Phe in BRAF, there is a large predicted shift in conformational equilibrium that results from destabilizing the N-terminal anchor (ΔΔ*T* ≈ −2). This appears to be a consequence of mutations abrogating favorable cross-strand contacts between the branched V120_HRD-2_ and unbranched A146_DFG+2_ or S146_DFG+2_ sidechains that are frequently present in wildtype STKs and display complementary packing between the C_β_ atom of A146_DFG+2_ or S146_DFG+2_ and the branched Cγ atom of I120_HRD-2_ or V120_HRD-2_ (Figure 4A). These complementary combinations of residue pairs are rare in TKs^15^, making their wildtype N-terminal anchors intrinsically less-stable relative to STKs. The Potts model predictions of these β6-β9 cross-strand interaction constraints were validated using FEP simulations; for CDK6 (an STK which contains A146_DFG+2_), mutating V120_HRD-2_ from a branched sidechain to Phe (V120F) abrogates favorable Cγ-C_β_ sidechain interactions with A146_DFG+2_, destabilizing the N-terminal anchor in the active conformation by nearly 4 kcal/mol relative to the inactive conformation (ΔΔ*G*_*reorg*_ = −3.7 ± 0.1 kcal/mol), consistent with the Potts threaded-energy predictions (ΔΔ*T* ≈ −2). As a further validation of this selection rule, for INSR (a TK bearing F120_HRD-2_ and T146_DFG+2_ as the wildtype) we find that substituting F120_HRD-2_ with a branched sidechain, F120I, stabilizes the active conformation as observed for STKs (ΔΔG_reorg_ = 2.5 ± 0.2 kcal/mol) only if the activation loop residue T146_DFG+2_ is first mutated to Ala (the STK-prevalent residue) to satisfy the interaction constraint described above, otherwise the Cγ atoms of wildtype T146_DFG+2_ and mutant F120I will clash, resulting in ΔΔ*G*_*reorg*_ ≈ −0.8 ± 0.2 kcal/mol.

#### 2.2. The RD-pocket is destabilized in the active conformation in TKs relative to STKs

The “RD-pocket” (Figure 4B) plays a distinctive functional role in STKs compared with TKs. Similar to what is seen for the N-terminal anchor in STKs, the RD-pocket in the active conformation of STKs directly interfaces with co-crystallized substrates, in contrast to TKs which bind peptides in a different binding mode^16^. The RD-pocket is a dynamically assembled and positively charged pocket formed by a cluster of basic and hydrophobic sidechains originating from the HRD-Arg (R123_HRD_), the activation loop N-terminus (145_DFG+1_ through 147_DFG+3_) and C-terminal anchor (159_APE-8_ – 161_APE-7_). The αC-helix also contributes basic residues to the RD-pocket in some kinases. For both TKs and STKs, the net-charge of this pocket places an additional contingency on the stability of the active conformation relative to inactive that is controlled by the biological environment: when Tyr, Ser, or Thr residues in the activation loop (153_DFG+9_ – 155_DFG+11_) are phosphorylated by another kinase, the acidic/phosphorylated sidechain buries itself into this pocket *in cis* and stabilizes the active conformation.^14^ For unphosphorylated kinases, this pocket can also be stabilized *in trans* by anions from the surrounding buffer as observed in our simulations (Figure 4B).

The apo, active conformation of STKs is stabilized by favorable packing between RD-pocket residues, namely those originating from the activation loop N-terminal and C-terminal residues (e.g., L145_DFG+1_ and V160_APE-7_ in CDK6: see Figure 4B). This is enabled in-part by residue 160_APE-7_ which, in STKs, has a small aliphatic sidechain, allowing it to pack tightly against the sidechain of residue 145_DFG+1_ and R123_HRD_, stabilizing the RD-pocket. In TKs however, residue 160_APE-7_ located in the C-terminal anchor is usually a bulky and/or positively charged residue (Lys or Arg) with the sidechain “flipped out” from the RD-pocket and exposed to solvent, e.g, K400 in Abl kinase numbering (PDG: 2G2I). Using FEP simulations of CDK6 to perform these substitutions, V160K_APE-7_ and V160R_APE-7_ which are frequently observed in TKs, we find an (unfavorable) increase in the free energy of the active conformation relative to the inactive conformation, resulting in ΔΔ*G*_*reorg*_ = −1.6 ± 0.1 and −2.6 ± 0.1 kcal/mol, respectively. This appears due to the structural decoupling of residue 160_APE-7_ from the RD-pocket, in concordance with the Potts calculated ΔΔ*T*s (ΔΔ*T* = −2.3 and −3.1, respectively). These results suggest the RD-pocket is relatively less stable when TKs are in the active conformation compared with STKs.

#### 2.3. The C-terminal anchor stabilizes the active conformation in STKs

The third structural motif in the activation loop, the C-terminal anchor, is directly involved with the binding of substrates to the active conformation *in trans* for both STKs and TKs^11,13^ by providing a complementary surface that stabilizes catalytic interactions between the substrate hydroxyl and the conserved HRD-Asp residue (D124_HRD_) in the catalytic loop.^37^ In STKs, this surface is formed in the active conformation by a conserved hydrogen bond between T162_APE-5_ (or S162_APE-5_) and the catalytic base K126_HRD+2_ (HRDxKxxN) (Figure 5A).

**Figure 5.**
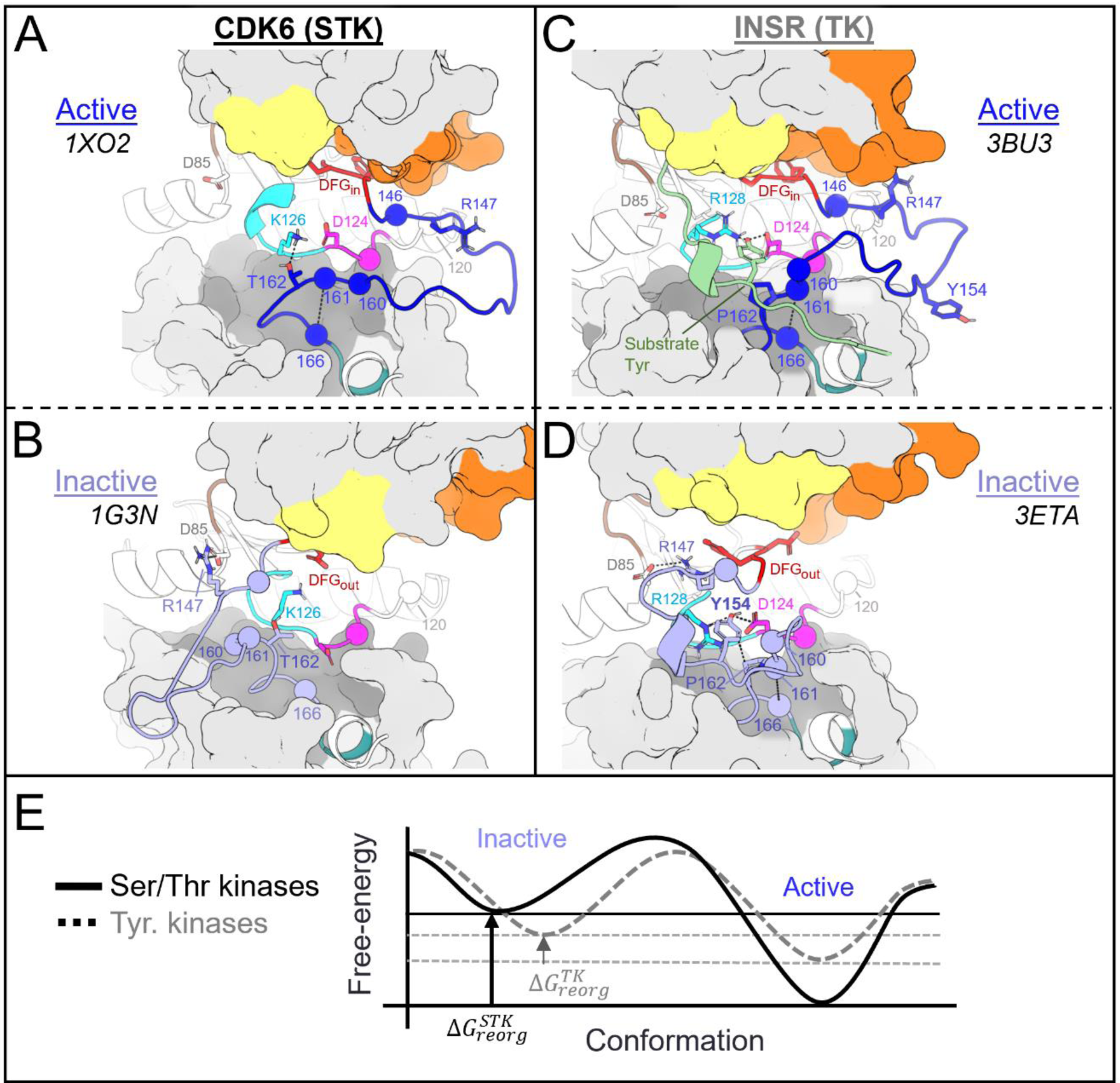
Divergent features of the active → inactive conformational change for TKs vs STKs. Key residues are displayed with α-carbon spheres and “sticks” representation. **(A)** Active conformation of CDK6 (PDB: 1XO2), an STK. In the active state the activation loop (dark blue) is extended and the C-terminal anchor is formed by a hydrogen bond between K126_HRD+2_ in the catalytic loop and T162_APE-5_ in the activation loop C-terminus. The activation loop C-terminus is also anchored in-place via the stacked residues 161_APE-7_ and 166_APE-6_. **(B)** Viewing the inactive DFG-out activation loop of CDK6 (PDB: 1G3N) – a large rotation about T162 can be seen which distorts the C-terminal anchor and breaks the contact between 161 and 166. **(C)** Viewing the active holoenzyme of INSR (PDB: 3BU3) with a peptide substrate bound to the active kinase in the C-lobe binding mode. **(D)** Viewing the inactive activation loop of INSR (PDB: 3ETA). Unphosphorylated Y154 in the activation loops of TKs (light blue) acts as a pseudo-substrate, forming the same interactions as the substrate phosphoacceptor in *C*. **(E)** Depiction of the active ↔ inactive landscape suggested by the Potts model and structural observations, for STKs (solid line) and TKs (dashed). The barrier heights are unknown and were drawn for the sake of illustration, while the relative depths of the active and inactive free energy basins were drawn descriptively, based on estimates of Δ*G*_*reorg*_ (see ref. 15) and ΔΔ*G*_*reorg*_.

When an STK is reorganized to the inactive DFG-out activation loop-folded conformation^15^ (Figure 5B), this hydrogen bond is broken as in CDK6 (active PDB 1XO2:B, inactive PDB 1G3N:A). This is due to a large torsion of the activation loop backbone about 162_APE-5_, an effect which can be seen by comparing Figure 5A and Figure 5B. We previously found that breaking this hydrogen bond is suggested by the Potts model to come at a large free-energy cost.^15^ Additionally, we found other sidechain interactions in the C-terminal anchor which are also broken upon reorganization of the activation loop in STKs; due to rotation of the backbone about 162_APE-5_, 161_APE-6_ is “swung” outwards in the inactive conformation (Figure 5B) and breaks its contact with 166_APE-1_ which incurs an additional energetic penalty according to the Potts model.^15^

#### 2.4. The C-terminal anchor stabilizes both the active and inactive conformation in TKs

##### Role of the C-terminal anchor in the active conformation of TKs

The C-terminal anchor of TKs is strikingly different from STKs; TKs have a conserved Pro at position 162_APE-5_ which rigidifies the local backbone and simultaneously provides a complementary surface in the active conformation for the binding of bulky Tyr sidechains *in trans* (Figure 5C). In TKs, the “catalytic base” is typically R128_HRD+4_ in place of the STK-conserved K126_HRD+2_. Additionally, in TKs the sidechains of contact-pair 161_APE-6_ and 166_APE-1_ in the C-terminal anchor form part of the “C-lobe” binding site located “underneath” the activation loop where substrate peptides are found co-crystallized with TKs in the active conformation.^16^

##### The distinct C-terminal anchors of TKs enable substrate mimicry in the inactive conformation

Unlike STKs, the C-terminal anchors of experimentally solved TKs in the inactive conformation are generally seen intact while the activation loop N-terminus and middle-region are “folded over” (Figure 5D), likely due to the enhanced rigidity of the C-terminal anchor in TKs. This allows the flexible middle-region of the activation loops to fold onto the C-terminal anchor *in cis* as a pseudosubstrate. The flexible middle-region of TK activation loops typically contain one or more Tyr residues, e.g., Y154_DFG+10_ which, in the inactive conformation, stacks onto P162_APE-5_ in the C-terminal anchor and interacts with the kinase’s own catalytic machinery.^17^ For inactive INSR structures (e.g., PDB: 3ETA), *cis* pseudo substrate interactions are identified by hydrogen bonds between the Y154_DFG+10_ hydroxyl and D124_HRD_ and R128_HRD+4_ in the catalytic loop (Figure 5D). The stability of these interactions was corroborated at the sequence level using threaded energies from the Potts model which predict favorable interactions between Y154_DFG+10_, the TK catalytic base (R128_HRD+4_), and P162_APE-5_ in the C-terminal anchor. As described below, the sequence-based predictions were confirmed at the structural level using FEP simulations to mutate Y154_DFG+10_ to residues seen in STKs.

STK activation loops are rarely observed with Tyr at the 154_DFG+10_ position and instead have a variable residue, Leu being the most frequent which appears in nearly 10% of STKs and only 3% of TKs. By contrast, Y154_DFG+10_ is highly prevalent in the activation loops of TKs (∼30%). The mutation Y154L in INSR was predicted by the Potts model to result in one of the largest possible single-mutant shifts in stability away from the inactive conformation and towards the active conformation. This predicted effect was confirmed by performing FEP simulations in both the active and inactive conformations, resulting in ΔΔ*G*_*reorg*_ = 3.3 ± 0.5 kcal/mol which is consistent with the Potts-calculated ΔΔ*T* = 1.1 (Figure 5E). A large fraction of this free-energy penalty to the inactive state may be due to the elimination of pseudo-substrate hydrogen bonds with D124_HRD_ and R128_HRD+4_, as suggested by the result of mutating Y154_DFG+10_ to Phe (Y1162F in INSR numbering). Similar to Y154L, Y154F tilts the free energy balance of the active and inactive basins in favor of the active basin, consistent with experiments that report an increase in basal activity for this mutant^17,38^. These results validate the proposed regulatory role^17^ of unphosphorylated activation loop Tyr for optimizing the stability of the inactive “substrate mimetic” conformation commonly seen in x-ray diffraction^23^ and NMR structures^27^ of TKs.

#### 2.5. Mutating the regulatory spine alters the stability of the active conformation relative to inactive

The regulatory spine or “R-spine” is a structurally conserved column of (typically) nonpolar, stacked sidechains located inside the kinase hydrophobic core that connects the N-lobe and C-lobe when the kinase is active^45,46^ and is thought to orchestrate correlated dynamics of the kinase domain which are required for catalysis.^47,48^ The R-spine is disassembled upon displacement of the DFG motif from “DFG-in” to “DFG-out”, or the αC-helix from “αC-in” to “αC-out”, and in this way the R-spine can be stabilized or destabilized by allosteric signals from the αC-helix and activation loop. The R-spine itself is structurally conserved throughout eukaryotic kinase families and plays a common functional role in both TKs and STKs. Our results from the Potts model and free energy simulations described next confirm the regulatory role of this structural motif in both kinase families.

##### Mutations of the αC-helix and DFG motif have consistent effects in the Potts model and FEP

In the active conformation the DFG motif participates in the R-spine via F143_DFG_ located at the “base” of the R-spine stack. Moving “up” the spine, the next residue belongs to the αC-helix located at position 53 in our MSA numbering, and it is usually a bulky sidechain (e.g., M53_αC_ or L53_αC_) that packs against the aromatic ring of F143_DFG_ in the active conformation. Several mutations of residues located in and around the R-spine were identified in our Potts mutational scan to result in large shifts in conformational preference between active and inactive: namely residues F143_DFG_ and M53_αC_ in INSR, and L53_αC_ in BRAF. We calculated ΔΔ*T* and ΔΔ*G*_*reorg*_for these mutations, the results for which are presented in Figure 3C and Table S3.

##### Results for mutating the DFG motif imply distinct conformational landscapes of DLG kinases

“DLG” is the most commonly observed variation of the DFG motif, seen in ∼10% of wildtype STKs and TKs, although its functional purpose is not well understood. The mutation “DFG to DLG” (F143L_DFG_) was predicted by the Potts model and confirmed with FEP simulations to have an activating effect in BRAF (ΔΔ*G*_*reorg*_ = 0.9 ± 0.4) and a larger inactivating effect in INSR (ΔΔ*G*_*reorg*_ = −2.9 ± 0.4 kcal/mol) whereby the active/assembled R-spine is stabilized in the former and destabilized in the latter. Analysis of Potts threaded energies in the wildtype and mutant (F143L) sequences provides a structural rationale for the differential effect of this mutation in INSR compared with BRAF in our FEP simulations (see Figure S2), which suggests the effect of the DLG substitution depends on the identity of residues surrounding the R-spine in the C-lobe. For INSR in the active DFG-in basin, Potts threaded energies predict favorable interactions between wildtype F143_DFG_ and residue 56_αC_ at the C-terminus of the αC-helix. Structurally, INSR has a bulky hydrophobic sidechain at this position (F56_αC_) which packs closely with F143_DFG_ in the R-spine (Figure S2), and the F143L_DFG_ mutation is predicted by the Potts model to weaken this interaction, thereby destabilizing the active conformation. In contrast, BRAF has a small Thr sidechain (T56_αC_) which does not interact with F143_DFG_ as closely, consistent with the Potts threaded energy for this interaction in the active state, which is very weak, and therefore the F143L_DFG_ mutation destabilizes the active conformation of INSR but not BRAF.

## Discussion

### 3. On the evolution of functional regulatory conformations in TKs and STKs

For TKs, the conformational free-energy basin corresponding to the inactive state is, due to features of the catalytic domain sequence that distinguish TKs from STKs, of similar depth to the active basin (approximately 1 kcal/mol relative free energy difference favoring the active state for TKs, on-average^15^) as illustrated in Figure 5E. We suggest how this bias may have evolved in tandem with tighter and more complex modes of regulation required for high fidelity intercellular phosphotyrosine signaling during the evolution of metazoan multicellularity.

Our analysis of mutational effects on kinase conformational landscapes, using the Potts model and free energy simulations suggest that, in the absence of extrinsic signals (such as phosphorylation on Y154_DFG+10_, protein-protein interactions, and domain assemblies^27,49,50^) the intrinsic propensities of TK catalytic domains for the active → inactive reorganization evolved with (A) structural features of the active conformation that differ substantially from STKs to shape a distinct peptide binding surface, and (B) the ability for TK activation loops to closely mimic *in cis* the binding mode adopted by TK-bound peptide substrates, which regulates formation of the holoenzyme and simultaneously shields the activation loop Y154_DFG+10_ from activating trans-phosphorylation by sequestering it within the kinase’s own active site as a pseudo substrate.

#### 3.1. The evolution of a distinct substrate binding mode in TKs explains the altered stability of the active, apo state in TKs

The three active-state motifs we identified to control the active → inactive conformational bias in TKs and STKs have a second functional role: they are involved in the binding of substrate peptides *in trans* which bind in different orientations in PDB co-crystal structures of TKs compared with distantly related STKs^16^ (Figure 6A). This leads us to hypothesize that the evolutionary divergence in substrate binding functionality that distinguishes holo TKs from STKs has also altered the stability of the active, apo conformation. The substrate binding surface occupied by peptides in co-crystal structures of holo TKs involves residues in the C-lobe and the activation loop C-terminus in the active state with the peptide bound underneath the activation loop, while STKs are co-crystallized with substrates occupying the “cleft” between the N-lobe and C-lobe and with the peptide resting “on top” of the activation loop (Figure 6B). The functionalization of the C-lobe binding mode and decommissioning of the cleft binding mode during the evolution STKs to TKs is reflected by sequence changes in the N-terminal anchor, RD-pocket and activation loop C-terminus in the active state, each of which structurally participates in the intermolecular binding surfaces. In this context, each of these structural motifs play a dual functional role, interfacing with peptide substrates on one hand and controlling the stability of the active conformation on the other^13^. This suggests the evolution of structural features driving TK substrate binding has reshaped the free-energy landscape, resulting in a shallower active basin compared with STKs.

**Figure 6.**
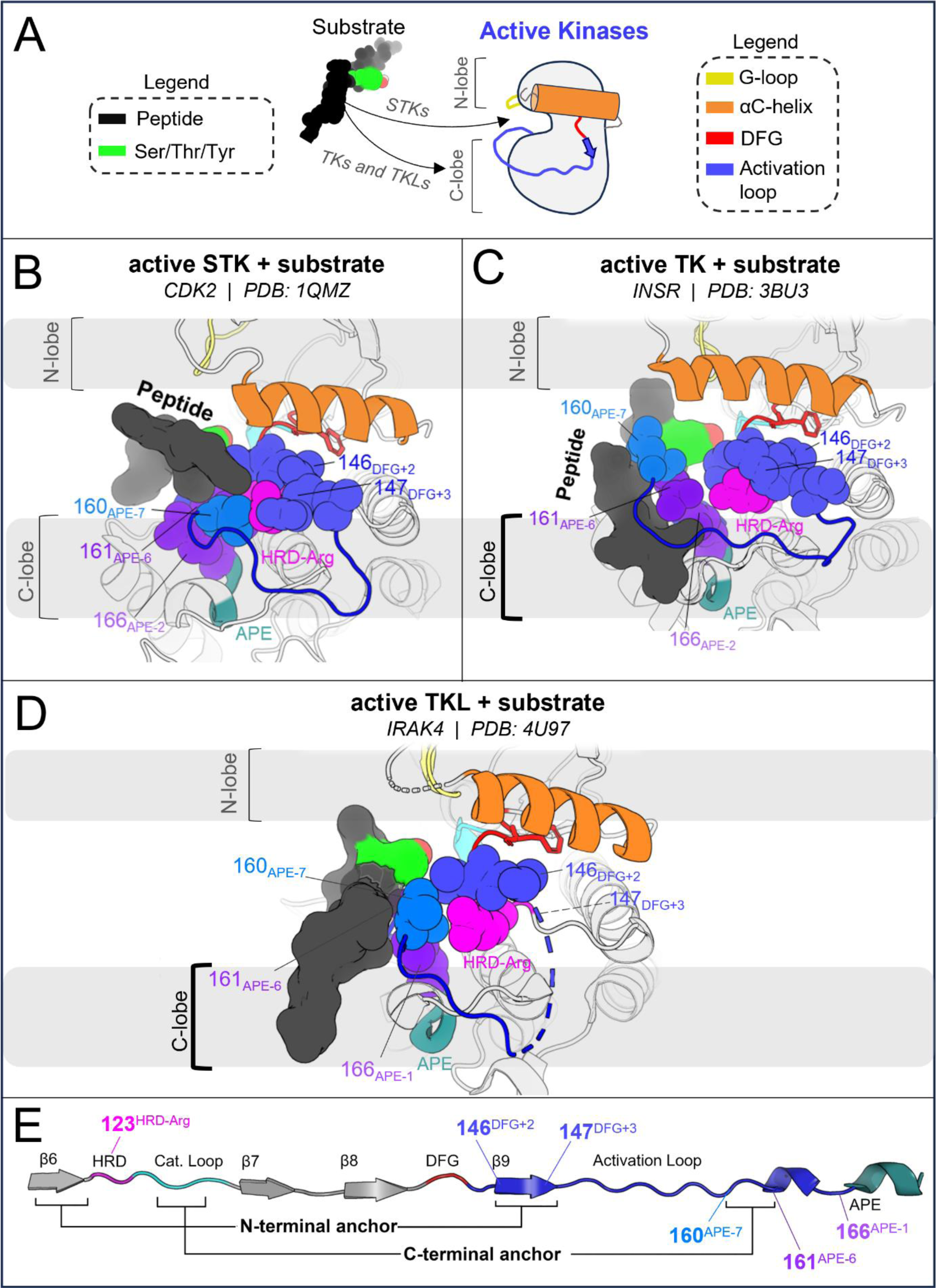
Structural differences between active and substrate-bound TK and STK co-crystal structures, where the TKL autophosphorylation structure of IRAK4 displays a similar substrate binding mode as TKs. (**A**) Cartoon illustration of substrate binding to TKs and STKs in different orientations, showing the angle-of-view for the catalytic domain used throughout Figure 6. (**B**) The catalytic domain of CDK2 bound to substrate (PDB: 1QMZ) containing a Ser phosphoacceptor. The peptide is bound within the “cleft”^16^ between the N-lobe and C-lobe. (**C**) The catalytic domain of INSR, a TK, bound to a substrate peptide (PDB: 3BU3) in the “C-lobe binding mode”^16^. (**D**) Viewing the trans-autophosphorylation dimer structure of IRAK4, a TKL. The activation loop from the opposing “substrate monomer” is colored as a substrate (dark grey) with downstream residues of the substrate monomer hidden for clarity. The substrate activation loop is oriented against the surface of active IRAK4 in a C-lobe binding mode, similar to the binding mode of peptide substrates to TKs. (**E**) Primary structure diagram showing the location and coloring of different motifs, with the residues depicted in *B-D* explicitly labeled. Residues that interact with peptides in the STK holoenzyme (e.g., CDK2) are labeled above, while residues that interact with peptides in the TK holoenzyme (e.g., INSR) are labeled below.

In the C-terminal segment of TKs, Arg and Lys are commonly seen at position 160_APE-7_ where a “bulge” in the activation loop forms part of the C-lobe groove occupied by peptides in TK co-crystal structures (Figure 6C). Above we showed these residues destabilize the active conformation relative to inactive. Structurally, “TK-like” mutants of CDK6 (an STK), such as V160R_APE-7_ and V160K_APE-7_, are observed in MD simulations to have an activation loop C-terminal segment that is structurally decoupled from the RD-pocket, causing it to distend outwards (Figure 3B) like what is seen in TKs^13^. Additionally, the stacked sidechains of residues 161_APE-6_ and 166_APE-2_ in the C-terminal anchor tend to be bulkier in TKs compared with STKs^15^, and form the inner “wall” of the C-lobe groove that interfaces with peptides (Figure 6C).

STKs generally do not have this C-lobe groove; their holo form is instead characterized by a binding mode where the peptide is placed “on top” of the activation loop and nested within the “cleft” between the “small lobe” (N-lobe) and “large lobe” (C-lobe) as illustrated in Figure 6B.^16^ As shown above, mutating the N-terminal anchor residues 120_HRD-2_ and 146_DFG+2_ of STKs to residues seen in TKs (e.g., CDK6: V120F and BRAF: A146T) significantly destabilizes the active conformation according to MD free energy simulations, tilting the free-energy landscape towards the inactive conformation. Because the “cleft” binding mode places peptides in contact with the activation loop N-terminus in the active state^16^ (Figure 6B) we propose that the decreased stability of the N-terminal anchor in TKs relative to STKs is due to the decommissioning of the cleft binding mode in TKs over the course of their evolution and the loss of functional constraints associated with this binding surface that otherwise require the N-terminal anchor to be highly stable, e.g., as is seen for STK.

In support of this hypothesis, co-crystallographic evidence in the PDB suggests that the TK-specialized “C-lobe binding mode” may have originated from the TKL group (Figure 6D), an “evolutionary intermediate” group of STKs that share a recent common ancestor with TKs.^9^ Strikingly, the TKL IRAK4 displays a C-lobe binding mode in its trans-autophosphorylation dimer structure^51^ PDB: 4U97 (Figure 6D) where the target substrate is a Thr residue^51–53^ (T345 in IRAK numbering). Relative to CDK2 (an STK that binds peptides in the cleft), IRAK4 has an unstable N-terminal anchor in the active conformation according to the Potts model, similar to TKs (see Table S1). This is due to the presence of a bulky aromatic residue in the N-terminal anchor, H120_HRD-2_ (H307 in IRAK numbering, see Figure S3), which has an effect on the active conformation analogous to F120 seen in TKs (e.g., F1128 in INSR numbering). This is the only example to our knowledge of an STK co-crystallized with a substrate in this binding mode, and although caution should be exercised when interpreting limited crystallographic data, our structural observations and Potts threading calculations for IRAK4 summarized in Figure S3 and Table S1 suggest the functionalization of the C-lobe binding surface and evolutionary “decommissioning” of the cleft binding surface may have originated in TKLs, prior to the evolution of Tyr-specific features of the active site. In other words, the evolution of the C-lobe binding mode may have preceded the more “fine-grain”, atomically precise arrangement of active site residues evolved by TKs for binding Tyr sidechains *in trans*.

#### 3.2. Co-evolution of autoinhibitory substrate mimicry in cis and substrate binding modes in trans

We additionally suggest that the autoinhibitory conformation of the INSR activation loop^17^ against the C-lobe *in cis,* which mimics the *trans* binding mode of peptides against the C-lobe in the active conformation, evolved from STKs to TKs in tandem with new regulatory modes, with TKLs as an “evolutionary intermediate”^8,54^. Comparing the inactive activation loops of apo CDK6, BRAF, and INSR in Figure 7 provides a qualitative view of how the substrate mimetic inactive conformation of TKs might have been shaped over the course of evolution from STKs → TKLs → TKs to regulate the kinase-preferred binding mode of peptides and protein substrates *in trans*. We note that STKs (including TKLs) have few structures in the PDB which are simultaneously apo and have the activation loop in a fully resolved DFG-out conformation. STKs belonging to the CDK family evolved as part of the early mitotic machinery in eukaryotes^55^ in contrast to TKs and TKLs which diverged only recently^56^ and display similar substrate binding modes oriented against the C-lobe *in trans* (Figure 6C and Figure 6D), and in this light it is not surprising that BRAF (Figure 7D), in contrast to CDK6 (Figure 7B), adopts an inactive activation loop conformation that more-closely resembles what is seen in TKs such as INSR with the activation loop folded against the C-lobe surface (Figure 7C). The inactive loop of CDK6 does not interface with the C-lobe surface, as expected if substrate-mimetic interactions between the activation loop and the C-lobe surface lack a functional purpose in CDK6 and STKs more broadly, with the exception of TKLs as noted.

**Figure 7.**
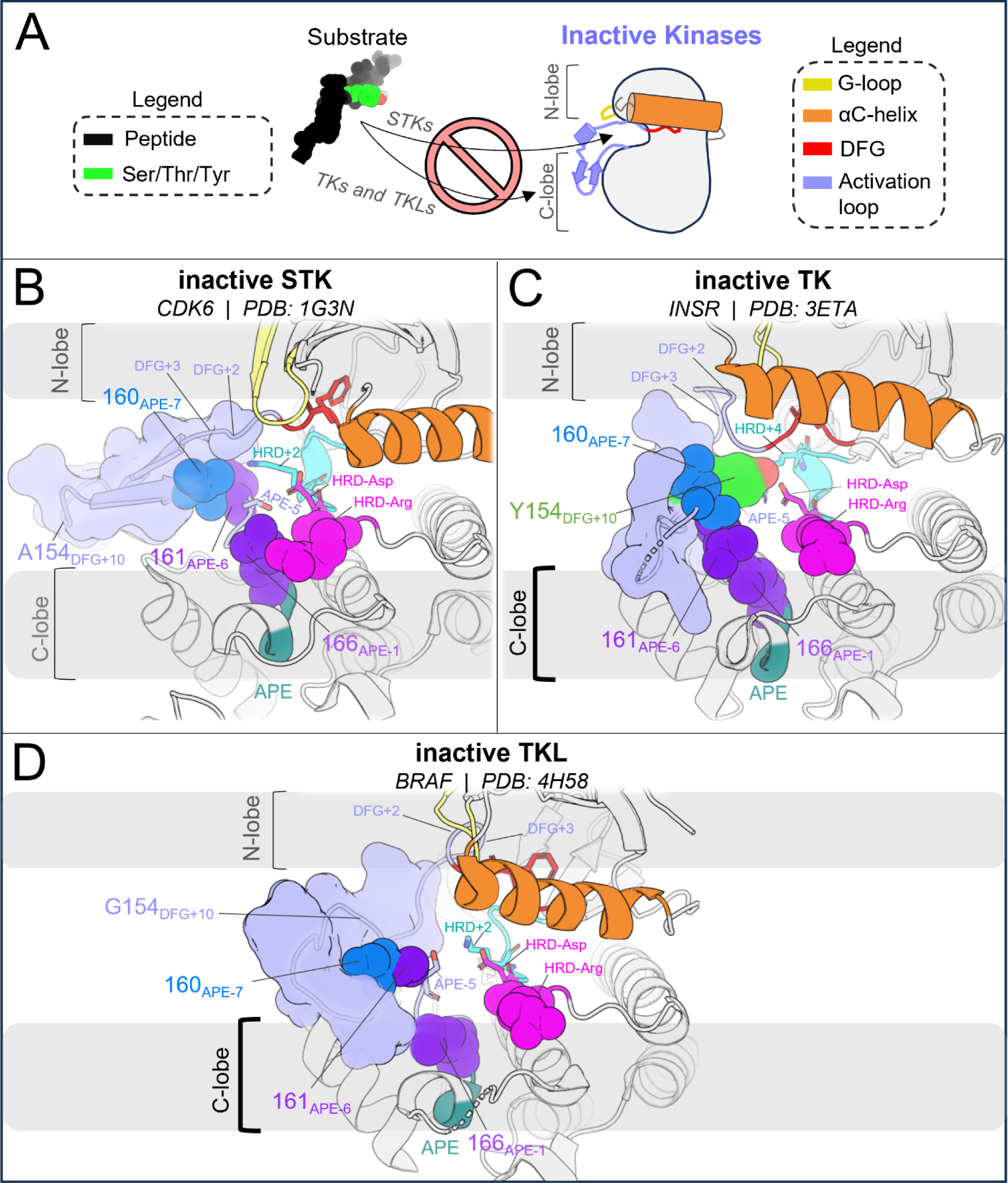
Substrate mimicry in the inactive conformation may have evolved in TKLs – a group of STKs that share a recent common ancestor with TKs. (**A**) Cartoon illustration of the inactive conformation which blocks substrates from binding, using the same angle-of-view as Figure 6. (**B**) Inactive CDK6 (PDB: 1G3N chain A), where the activation loop is highly reorganized relative to the active state but does not interact with the C-lobe surface where substrates bind to TKs. (**C**) Inactive INSR (PDB: 3ETA chain A) with the activation loop reorganized such that the middle-region *in cis* mimics a substrate with the activation loop Tyr (green) bound to the active site and downstream residues “blocking off” the C-lobe binding surface. (**D**) Inactive BRAF (PDB: 4H58 chain C) with an autoinhibitory activation loop that interacts with the C-lobe surface *in cis,* similar to INSR. However, by contrast BRAF lacks the “atomically precise” substrate mimicry involving a Tyr from the activation loop.

### 4. Concluding Remarks

In conclusion, our results and observations suggest the distinctive tilt of the TK free-energy landscape favoring the **DFG-out, activation loop-folded** inactive state relative to STKs is tied to features of their substrate binding functionality that distinguish TKs from STKs – the large-scale functionalization of the C-lobe surface for binding substrate peptides and, at a finer scale, the re-structuring of the active site in TKs to complement Tyr sidechains bound *in trans*. This physical restructuring of the active conformational basin has shifted the intrinsic free-energy landscape of apo TKs so that the balance more readily favors the inactive state for TKs compared with STKs. In addition to these differences in the stability of the active state arising from differences between peptide substrate binding modes of TKs and STKs, in the apo inactive state the “folded” activation loop of TKs behaves as a pseudo substrate and is stabilized by Tyr-specific features of the catalytic domain.

Integrating the analysis of the effects of mutations on the relative stability of active (DFG-in, activation loop extended) vs. inactive (DFG-out, activation loop-folded) kinase conformations using a sequence-based approach (Equation 3) with a structure-based approach (Equation 4) requires that both methods are sensitive enough to distinguish physicochemical differences between the two folded protein environments. Results from both computational approaches described by Equation (3) and Equation (4) are consistent, suggesting that the Potts statistical energy model and structure-based molecular dynamics free energy methods are presently at the level of accuracy where they are capable of capturing effects of mutations on the relative stability of two folded protein conformations which are both involved in function that individual proteins move between.

There is a strong correspondence between the changes in the conformational landscape induced by mutations, estimated using molecular dynamics simulations to construct alchemical thermodynamic cycles, and the results obtained using the Potts statistical energy model to thread sequences onto the corresponding structures in the active and inactive conformational free energy basins. We observe from both approaches that TKs have a less-stable active (DFG-in activation loop extended) conformational basin and a more-stable inactive (DFG-out activation loop-folded) conformational basin as compared with STKs (Figure 5E). A key benefit of the sequence-based model is the ability to decompose the statistical energy into pairwise interaction contributions, which can provide insights into the molecular basis of evolutionarily diverged conformational phenotypes, especially the role of the N-terminal and C-terminal anchors and the RD-pocket in stabilizing the active conformation of STKs. In contrast, the destabilization of the N-terminal anchor and RD-pocket in active-state TKs, together with the stable autoinhibitory interactions acquired by the TK clade involving Tyr-substrate mimetic interactions from the folded activation loop *in cis*, are the major determinants that distinguish active vs. inactive conformational states of TKs from STKs. As illustrated in Figure 7, a notable feature of the DFG-out activation loop-folded conformation for TKs compared with STKs is the ability to occupy a functional surface of the kinase that is otherwise “intended” for binding peptides and Tyr substrates. These structural observations, combined with the fact that many of the mutations with the strongest effects on the conformational landscape are also responsible for the evolutionary divergence in substrate specificity of TKs from STKs, support the proposal that the DFG-out conformation with the activation loop folded is a functional state that has been stabilized in TKs over the course of their evolution from STKs via the accumulation of residue substitutions in the activation loop and catalytic loop. This distinctive property of TKs likely enabled the evolution of tighter regulatory controls required for high-fidelity phosphotyrosine signaling.

## Methods

### System preparation and molecular dynamics simulations

Protein structures were prepared for MD simulations using Schrödinger suite 2023-2 version. For each of the three kinases studied, two x-ray crystallographic structures were downloaded from the PDB: CDK6 (active – 1XO2 chain B, inactive – 1G3N chain A), BRAF (active – 4MNE chain B, inactive – 4H58 chain C), and INSR (active – 3BU3 chain A, inactive – 3ETA chain A). All ligands, bound peptides and cofactors were stripped from the starting crystal structures. Crystallographic mutations were reverted back to wildtype and missing sidechains were added using Prime. Hydrogen atoms were added to the protein structures, and small loops were modeled using the Protein Crosslink tool. To determine ionization states, estimates of sidechain pKa values for titratable residues were obtained using PropKa. All crystallographic water molecules were retained. Restrained energy minimization was performed using the OPLS4 force field. For both the active and inactive DFG-out structures, we ensured the HRD histidine was singly protonated at the epsilon position, allowing it to form a hydrogen bond with the backbone of DFG-1 residue. We also ensured the conserved regulatory spine Asp was deprotonated and forms bidentate hydrogen bonds with the HRD-His and HRD-Arg backbone.

To relieve structural artifacts induced by the crystal packing environment, 100 ns molecular dynamics (MD) simulations were run in explicit solvent (TIP3P) with the Desmond simulation engine at constant temperature and volume, using the OPLS4 force field.^57^ An orthorhombic box geometry was used for periodic boundary construction, with minimum dimensions satisfying a 10 Å buffer between the protein and box boundaries. The simulations were run in the NVT ensemble (constant volume, temperature, and number of particles) with a buffer concentration of 0.15 M NaCl and additional ions as necessary to neutralize the system. At approximately 50 ns, a simulation frame was saved for further processing in preparation for FEP simulations. The first 50ns of the equilibration run were then discarded, and the last 50ns of the overall 100ns trajectory were used for Potts threaded energy calculations (see below).

### Free-energy Perturbation (FEP+/REST2) protocol and thermodynamic cycle for the calculation of *ΔΔG_reorg_*

From the MD frame extracted at 50ns, only the protein and water molecules within 5 Å of the protein were retained in preparation for FEP simulations. Ions from the surrounding salt buffer that were seen to interact with the protein and remained bound for a significant fraction of the simulation time were also retained. Ionizable residues were then subjected to PropKa calculations to determine ionization states, followed by another round of energy minimization. For mutation free-energy simulations, the FEP+/REST2 method^58,59^ implemented in the Desmond MD engine from Schrodinger Suite 2023-2 was used. The sidechain conformation of the mutant residue was initialized using Prime as part of the FEP+ default pipeline. For mutations involving changes in ionization state, 24 simulation replicas were initialized in 24 different lambda windows along an alchemical path separating the wildtype (*λ* = 0) and mutant (*λ* = 1) states. For all other mutation types, a total of 16 lambda windows were used. Replicas were simulated for 10 ns at constant volume and temperature with GCMC (grand canonical Monte-Carlo) sampling of water molecules between the bulk solvent and protein interior. Some mutations were run significantly longer to achieve convergence (up to 100ns, e.g., V160R). We ensured that both the active and inactive legs of the thermodynamic cycle were simulated for the same amount of time to avoid biasing the convergence of one leg over the other. Solute tempering was applied to residues within a sidechain heavy-atom distance of 5Å from the double mutant pair. This typically includes around 20 residues or hundreds of atoms. The solute tempering scheme applied to the double mutants was retained for all constituent single mutations and cycle closure calculations (described below). The replica exchange simulations were analyzed using the Benett Acceptance Ratio (BAR) implemented in the FEP+ program to calculate Δ*G*s for the alchemical legs. The difference between the physical legs, ΔΔ*G*_*reorg*_, was calculated by taking the difference Δ*G*(*B*) − Δ*G*(*A*) where *B* and *A* are the inactive and active basins, respectively.

### Kinase Potts model threaded energy calculation

The sequence-based calculations in this work were performed using the kinase Potts model from our previous paper^15^, where we refer the reader for more information. In brief, the q-state Potts model is a discrete statistical energy Hamiltonian that models co-evolutionary couplings *J* between all 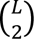 pairs of positions, where *L* = 259 is the number of aligned positions in our kinase MSA, and all possible combinations of the *q* = 21 amino acids letters (including a gap character) at those positions.

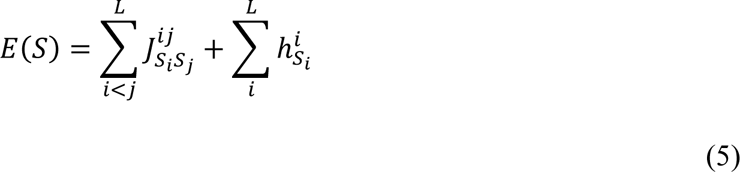

The total statistical energy *E* of sequence *S* determines its probability of being observed in the MSA relative to all other sequences which has the form of a Boltzmann distribution *P*(*S*) = 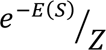 where *Z* is the partition function. The Mi3 algorithm uses MCMC to sample sequences from this distribution, starting from an initial “site-independent” guess for the Hamiltonian, and using Zwanzig reweighting and quasi-Newton optimization to iteratively converge on a set of coupling parameters that generate an ensemble of sequences with pairwise correlations precisely matching what is seen in the training MSA. The self-interaction field terms ℎ do not need to be inferred during model training, as the Potts model can be determined in a “fieldless” gauge where all the self-interaction energies are present implicitly as a constant shift in the coupling parameters. The fieldless Hamiltonian is then transformed to a “zero-mean” gauge shown in Equation 5 where the average coupling energies are set to zero and the remaining energy is distributed into the field terms. We do the threading calculation in this gauge, where pairs of residues with 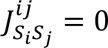 can be interpreted as uncoupled.^39^ The field terms do not contribute to the threading calculation described below.

The threaded energy of a sequence over a particular structure *n* from conformational ensemble *A* is calculated from the set of residue-residue contacts observed in that structure, by only summing Potts couplings of residue pairs that coincide with a nearest-sidechain heavy atom distance of < 6Å, represented by the delta function in the following formula:

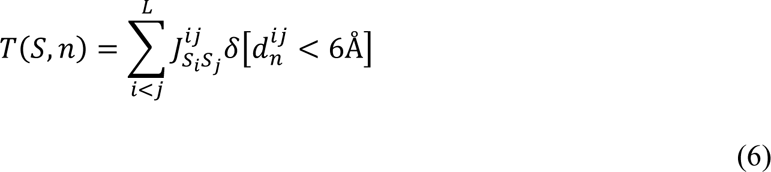

The average threaded energy of sequence *S* in basin *A* can be determined by substituting the delta function for an average – contact frequency 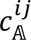:

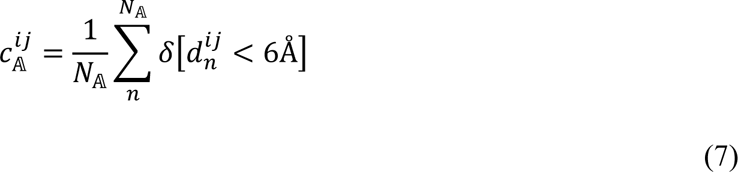

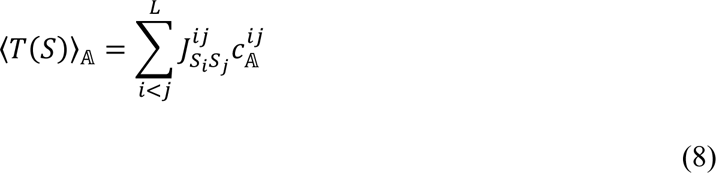

The average Potts threaded-energy difference between two basins, *A* and *B* is defined as Δ*T*(*S*),

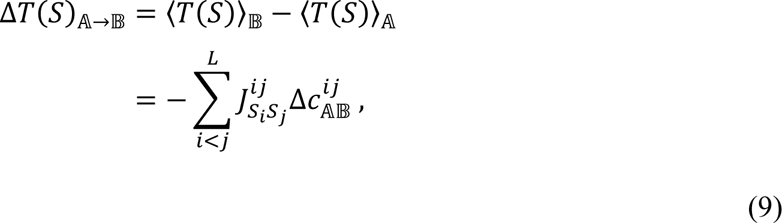

where the difference in contact frequencies is defined as 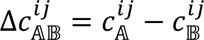.

The conformational ensembles *A* and *B* are defined as active DFG-in and inactive DFG-out, respectively, and the corresponding contact frequencies *c*_*A*_and *c*_*B*_are calculated using equally spaced 100 ps frames from the 50 ns trajectory of the threaded kinase, or PDB structures of homologous kinases as demonstrated for INSR. Because of the large availability of DFG-out structures for TKs in the PDB, we were able to use those in place of the MD ensemble for threading the INSR sequence consistent with the approach in our previous threading studies ^15,39^. Missing residues in PDB structures were accounted for by excluding their contribution to the contact frequencies (Equation 7).

### Calculation of threaded energy differences (ΔΔT) and construction of Potts mutation libraries

In order to scan for mutations in the Potts model that result in the largest shifts in threaded energy towards the active basin *A* or inactive basin *B* which can be confidently validated with FEP simulations, we limited our scan to all residues besides Gly, Pro, Trp, His, or Cys which might raise modelling complications due to large changes in backbone flexibility (Gly or Pro), bulky substitutions (Trp), or ambiguity regarding tautomer or ionization states and the potential for disulfide bridges (His or Cys). Positions in the sequence with any of these residues as a wildtype were omitted. Additionally, we limited the scan to positions that participate in at-least one interaction where the magnitude of their contact difference 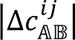 is greater than 0.8. This has the effect of limiting the mutation search to positions that undergo significant changes in their physicochemical environment. For these positions and their pairs, all possible single and double mutations (limited as described above) were performed, totaling between 10^4^ and 10^5^ ΔΔ*T* calculations for each kinase, i.e., 64,798 mutations for INSR, 366,336 for BRAF, and 808,960 for CDK6. The top three mutations were taken from the left tail (ΔΔ*T* ≪ 0 in Figure 3A) indicating a shift in stability that favors the inactive basin, and three additional mutations were taken from the right tail (ΔΔ*T* ≫ 0) which correspond to shifts in stability favoring the active basin, totaling six mutations. All six mutations for each of the three kinases correspond to changing pairs of residues, i.e., double mutations, resulting in 18 double-mutations made in both the active and inactive conformational free-energy basins (18 double-point mutations × 2 basins = 36 simulations).

### Single-point mutations and FEP double-mutant cycle closure calculations

In addition to the 18 double-mutations, 36 single-point mutations were performed with FEP in the active and inactive basins (36 single-point mutations × 2 basins = 72 simulations) for comparison with Potts ΔΔ*T*s which allowed us to assess cycle closure for the resulting double mutant cycles. This resulted in (18 + 36) × 2 = 108 FEP simulations in the active and inactive basins as shown in Table S3. Cycle closure was assessed by performing 36 additional FEP simulations, where the single mutants served as the initial state and the double mutants as the final state. In brief: there are three thermodynamically equivalent paths for arriving at the double mutant, by mutating residue *i* before residue *j*, *j* before *i*, or *i* and *j* simultaneously. Ideally, the calculated double mutant ΔΔ*G*_*reorg*_would be identical for all three paths resulting in a fully closed thermodynamic cycle. ΔΔ*G*_*reorg*_was plotted in Figure 3C after averaging the three different paths for the double mutation. The error bars in Figure 3C were calculated by taking the standard deviation of the three different paths, an estimate of the error in ΔΔ*G*_*reorg*_ due to hysteresis. There are also three paths to calculate ΔΔ*G*_*reorg*_for a single mutant using the double mutant cycle, which we used to report the average and standard deviation in Figure 3C and Table S3.

### Structural classification of tyrosine kinases

For INSR, the large availability of homologous proteins crystallized in both the DFG-out conformation with the activation loop “folded” and the active conformation with the activation loop “extended” allowed us to sample basins *A* and *B* from the PDB, similar to our previous work involving Potts threading. The KinCoRe database was used for defining the two ensembles; briefly, the spatial and dihedral classifiers calculated by Modi and Dunbrack ^21^ were used to define the active state as “DFG-in” and “BLAminus” respectively, where the spatial DFG conformation refers to the spatial location of the “Asp-Phe-Gly” motif at the beginning of the activation loop, and “BLAminus” refers to locations on the Ramachandran map for the residue immediately before the DFG motif as well as the Asp and Phe residues, i.e., “beta strand – left handed helix – right handed helix”, and a gauche-minus *χ*_1_rotamer state of the Phe sidechain. The inactive state is defined as “DFG-out” and “BBAminus” where the Asp of the DFG motif is flipped out of the ATP pocket and replaced with the adjacent Phe sidechain which occupies the ATP pocket underneath the glycine-rich loop (also called the P-loop), and the dihedral state of the XDF residues are “beta strand – beta strand – right-handed alpha helix” with the Phe sidechain in a gauche-minus state which has a strong correspondence with the activation loop in a “folded” conformation that deviates from the active “extended” conformation by ∼18Å. We imposed additional constraints on the active and inactive conformations: (A) both the active and inactive states display the αC helix is an “in” state, where the β3-Lys – αC-Glu salt bridge is formed. (B) both states display an “extended” glycine-rich loop, defined by the presence of a hydrogen bond between the backbone carboxyl of the first conserved glycine at the beginning of the loop (G1003 in PDB: 3BU3) and the backbone nitrogen of the conserved Val at the end of the loop in the β2-strand (V1010 in 3BU3) with a distance less than 3.5 Å. Our definition of “glycine-rich loop extended” also includes a constraint on the minimum distance (6.5 Å) between the G1003 carboxyl and the S1006 amino nitrogen (the residue immediately before the conserved Phe/Tyr) to avoid including structures with “kinked” glycine-rich loops^60^, which appears to be a feature of certain drug-bound kinases (e.g., PDB: 1IEP) but not INSR (PDB: 3ETA). This resulted in *N*_*A*_ = 437 PDB chains for TKs in the active basin and *N*_*B*_ = 157 chains for TKs in the inactive basin, used for the calculation of contact frequencies via Equation 7.

### Multiple sequence alignment and calculation of residue frequencies

Using the HHblits algorithm^61^ and starting from a hidden Markov model seed of the protein kinase superfamily from Pfam (PF00069) and filtering out sequences missing the DFG-Asp or which contain many gaps, an MSA containing 236,572 non-duplicate catalytic domain sequences were obtained. TKs and STKs were classified according to sequence constraints in the catalytic loop and activation loop as done previously^15^, resulting in 10,345 and 210,862 raw TK and STK sequences, respectively. When down-weighting each sequence with the number of similar sequences in our MSA (within 40% identity), the effective counts are 1,069 and 22,893 sequences respectively, which is a less biased estimate of the sequence diversity. This down-weighting procedure was used in the calculation of residue-pair and single-residue frequencies (marginals) which are reported in the main text as percentages (%).

## Data Availability

The Potts model and FEP calculations plotted in Figure 3C and described in the main text (ΔΔ*T* and ΔΔ*G*_*reorg*_) can be found in Table S3 provided in the Supplementary Information.

## Acknowledgements

This research has been supported by National Institutes of Health grants, number R35 GM132090, and R01 AI177849 and by NIH Computer Equipment Grant (OD020095).

