## Supplementary Information for "Evolutionary sequence and structural basis for the distinct conformational landscapes of Tyr and Ser/Thr kinases"

**Competing Interest Statement:** The authors declare no competing interests.

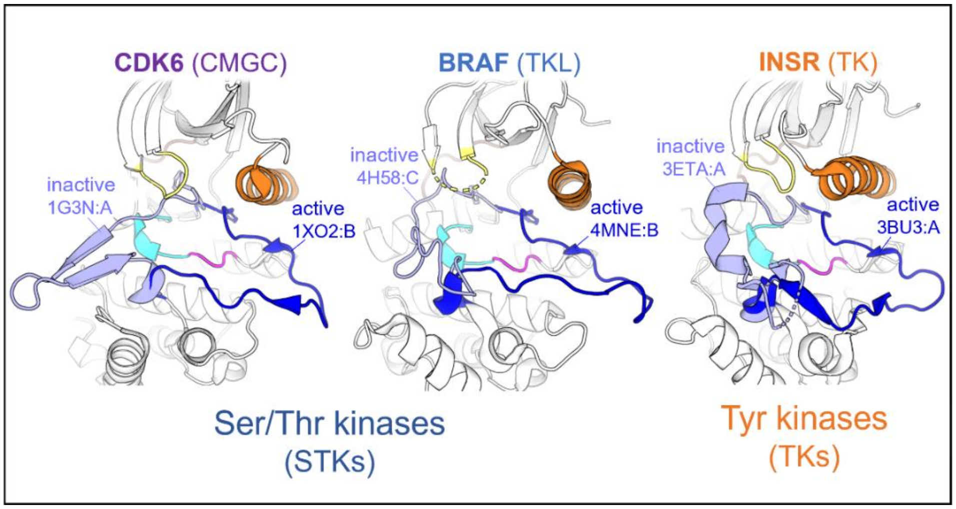


**Figure S1.** Overview of the three kinase targets and their active and inactive states, analyzed by Potts threading and molecular dynamics free energy simulations in this work.

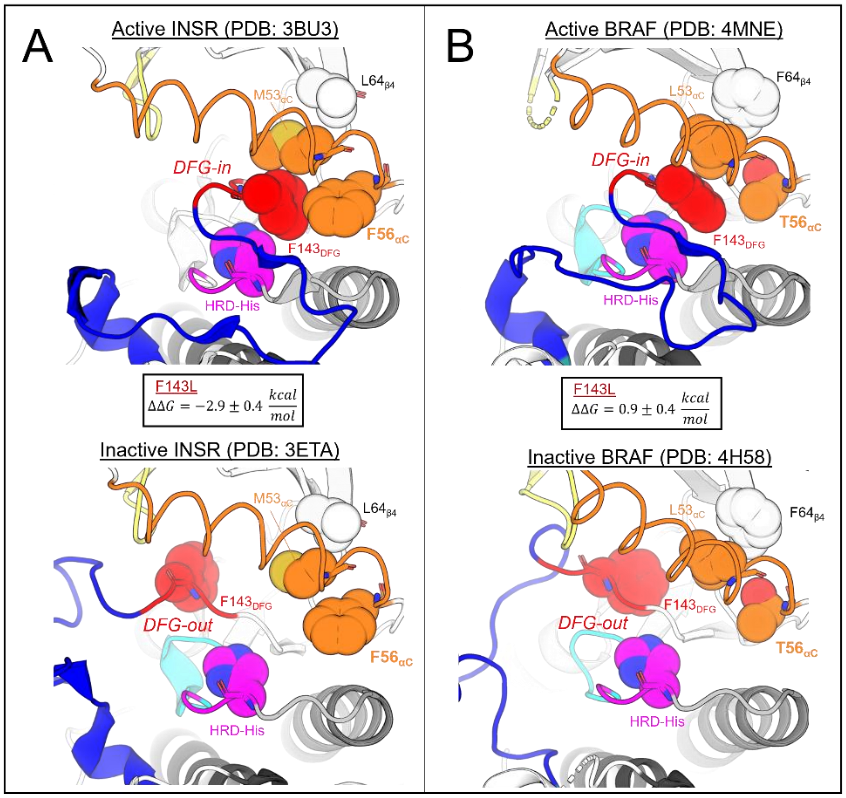


**Figure S2.** Potts threaded energies predict F143L to destabilize the R-spine of INSR, but not BRAF, consistent with FEP simulations (see Table S3 for data). The R-spine residue positions are labeled HRD-His, $143_{DFG}$, $53_{\alpha C}$ and $64_{\beta 4}$. Residue $56_{\alpha C}$ is located next to the R-spine in the αC-helix. $\Delta\Delta G_{reorg}$ for the $F143L_{DFG}$ mutation is reported in the inset. Potts threaded energies suggest that interactions between $F143_{DFG}$ and $56_{\alpha C}$ are weaker in BRAF compared with INSR. Consequently, the $F143L_{DFG}$ mutation penalizes the active-state stability of the INSR R-spine, in contrast to BRAF where the mutational effect has an opposite sign. (A) Active DFG-in vs inactive DFG-out conformation of INSR. (B) Active DFG-in vs inactive DFG-out conformation for BRAF.

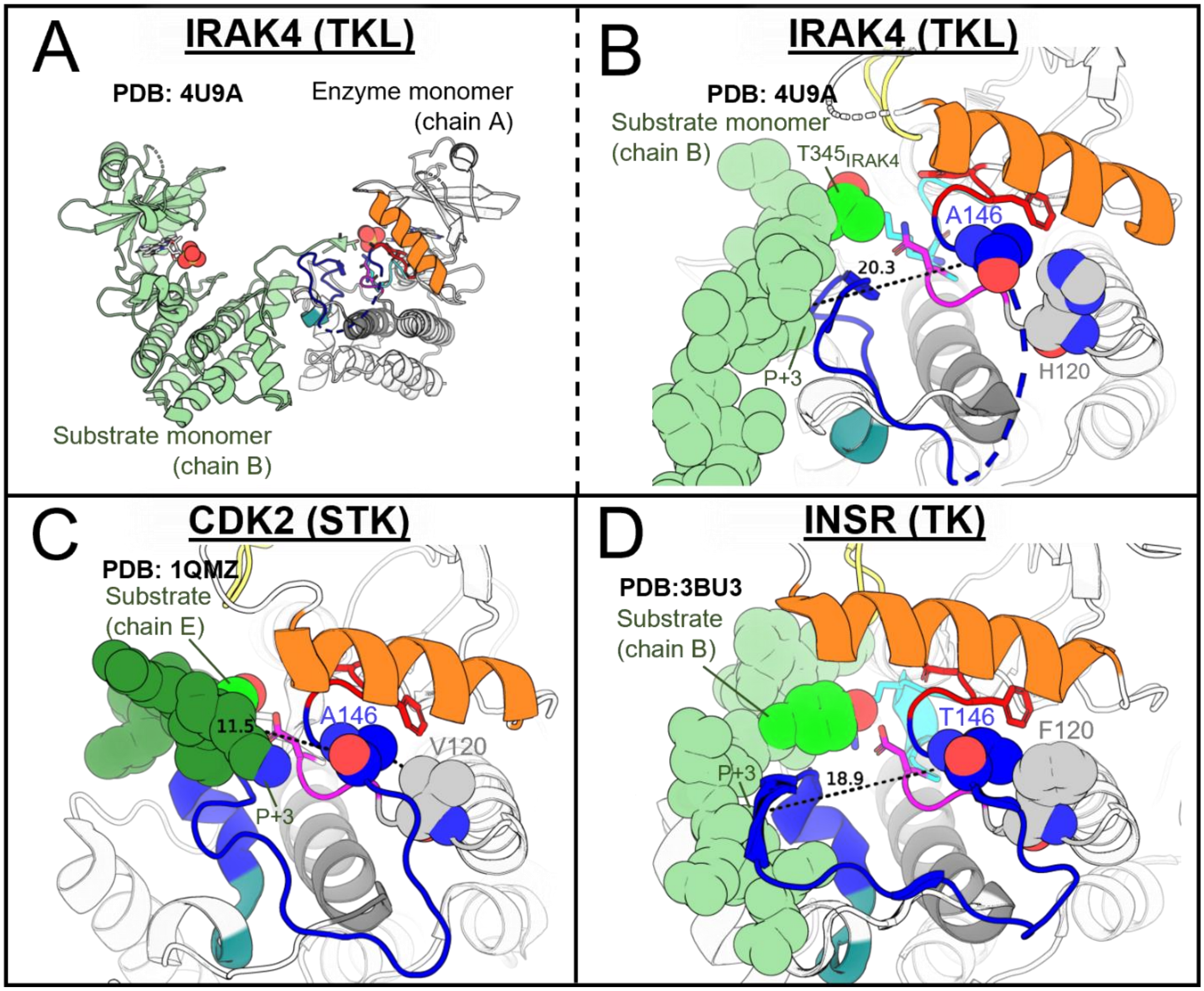


**Figure S3.** (**A**) Overview of the IRAK4 trans-phosphorylation dimer structure, PDB: 4U97. (**B**) Viewing the IRAK4 enzyme monomer and substrate activation loop from the opposing chain. Chain A represents the active enzyme monomer while a Thr residue (T345) from the activation loop of chain B serves as the substrate/phosphoacceptor. (**C**) CDK2 bound to a substrate peptide (PDB: 1QMZ) with the peptide oriented into the kinase cleft between the N-lobe and C-lobe (see Figure 6 for schematic). The peptide is bound nearby the DFG+2 residue (A146) in the activation loop N-terminal anchor. Relative to CDK2 (an STK), IRAK4 has an unstable N-terminal anchor according to the Potts model due to $H120_{HRD-2}$ (H307 in IRAK numbering), similar to the effect of F120 in TKs (e.g. INSR, see Figure 3). (**D**) INSR bound to a substrate peptide. INSR and many other TKs have a bulky aromatic residue at this position (F120) which has destabilizing effects on the activation loop N-terminus, consistent with evolutionary "decommissioning" of the cleft-oriented binding mode in TKs. Our Potts model calculations in Table S1 might suggest that H120 is present in IRAK4 for a similar reason – the STK-preferred "cleft" binding mode might be decommissioned in IRAK4, and possibly other TKLs, consistent with the substrate-bound structure in *A-B* and the nature of TKLs as "evolutionary intermediates" along the evolutionary path between STKs and TKs (Figure 2).

**Table S1.** Potts model results for each of the three kinases from Figure S3. Smaller values of $\Delta T$ reflect destabilization of the extended/active conformation of the activation loop relative to the inactive conformation. Mutating the STK-prevalent V120 in CDK2 to the residue seen in IRAK4 (V120H) destabilizes N-terminal anchor according to the Potts model ($\Delta\Delta T \approx -1$), and the reverse mutation in IRAK4 (H120V) recovers some stability of the N-terminal anchor ($\Delta\Delta T \approx +1$) to make the kinase more "STK-like". This is consistent with the observation in Figure S2B that IRAK4 utilizes the TK-preferred peptide binding mode, where the peptide (in this case, the activation loop of a substrate monomer) is bound far away from the enzyme's activation loop N-terminus.

| Class | Kinase | HRD-2 (pos. 120) | Wildtype Potts ΔT | Mutation | ΔΔT |
|---|---|---|---|---|---|
| STK | CDK2 | Val | 5.3 | V120H | -1.2 |
| TK | INSR | Phe | 0.5 | F120H | 0.1 |
| TKL | IRAK4 | His | 2.5 | H120V | 0.89 |

**Table S2**. Alignment to PDB numbering conversion table.

| Alignment | CDK6 (1G3N:A) | BRAF (4H58:C) | INSR (3ETA:A) |
|---|---|---|---|
| 1 | - | - | - |
| 2 | Q12 | Q455 | K995 |
| 3 | Y13 | I456 | I996 |
| 4 | E14 | T457 | T997 |
| 5 | C15 | V458 | L998 |
| 6 | V16 | G459 | L999 |
| 7 | A17 | Q460 | R1000 |
| 8 | E18 | R461 | E1001 |
| 9 | I19 | I462 | L1002 |
| 10 | G20 | G463 | G1003 |
| 11 | E21 | S464 | Q1004 |
| 12 | G22 | G465 | G1005 |
| 13 | A23 | S466 | S1006 |
| 14 | Y24 | F467 | F1007 |
| 15 | G25 | G468 | G1008 |
| 16 | K26 | T469 | M1009 |
| 17 | V27 | V470 | V1010 |
| 18 | F28 | Y471 | Y1011 |
| 19 | K29 | K472 | E1012 |
| 20 | A30 | G473 | G1013 |
| 21 | R31 | K474 | N1014 |
| 22 | D32 | W475 | A1015 |

| Alignment | CDK6 (1G3N:A) | BRAF (4H58:C) | INSR (3ETA:A) |
|---|---|---|---|
| 23 | L33 | - | R1016 |
| 24 | N35 | - | E1022 |
| 25 | G36 | - | A1023 |
| 26 | G37 | H476 | E1024 |
| 27 | R38 | G477 | T1025 |
| 28 | F39 | D478 | R1026 |
| 29 | V40 | V479 | V1027 |
| 30 | A41 | A480 | A1028 |
| 31 | L42 | V481 | V1029 |
| 32 | K43 | K482 | K1030 |
| 33 | R44 | M483 | T1031 |
| 34 | V45 | L484 | V1032 |
| 35 | R46 | N485 | N1033 |
| 36 | V47 | V486 | E1034 |
| 37 | Q48 | T487 | S1035 |
| 38 | T49 | A488 | A1036 |
| 39 | G50 | P489 | S1037 |
| 40 | E52 | P491 | L1038 |
| 41 | G53 | Q492 | R1039 |
| 42 | M54 | Q493 | E1040 |
| 43 | P55 | L494 | R1041 |
| 44 | L56 | Q495 | I1042 |
| 45 | S57 | A496 | E1043 |
| 46 | T58 | F497 | F1044 |
| 47 | I59 | K498 | L1045 |
| 48 | R60 | N499 | N1046 |
| 49 | E61 | E500 | E1047 |
| 50 | V62 | V501 | A1048 |
| 51 | A63 | G502 | S1049 |
| 52 | V64 | V503 | V1050 |
| 53 | L65 | L504 | M1051 |
| 54 | R66 | R505 | K1052 |
| 55 | H67 | K506 | G1053 |
| 56 | L68 | T507 | F1054 |
| 57 | E72 | R508 | T1055 |
| 58 | H73 | H509 | C1056 |
| 59 | P74 | V510 | H1057 |
| 60 | N75 | N511 | H1058 |
| 61 | V76 | I512 | V1059 |
| 62 | V77 | L513 | V1060 |
| 63 | R78 | L514 | R1061 |
| 64 | L79 | F515 | L1062 |

| Alignment | CDK6 (1G3N:A) | BRAF (4H58:C) | INSR (3ETA:A) |
|---|---|---|---|
| 65 | F80 | M516 | L1063 |
| 66 | D81 | G517 | G1064 |
| 67 | V82 | Y518 | V1065 |
| 68 | C83 | S519 | V1066 |
| 69 | T84 | T520 | S1067 |
| 70 | V85 | K521 | K1068 |
| 71 | S86 | P522 | G1069 |
| 72 | R87 | - | Q1070 |
| 73 | K93 | Q523 | P1071 |
| 74 | L94 | L524 | T1072 |
| 75 | T95 | A525 | L1073 |
| 76 | L96 | I526 | V1074 |
| 77 | V97 | V527 | V1075 |
| 78 | F98 | T528 | M1076 |
| 79 | E99 | Q529 | E1077 |
| 80 | H100 | W530 | L1078 |
| 81 | V101 | C531 | M1079 |
| 82 | D102 | E532 | A1080 |
| 83 | - | G533 | H1081 |
| 84 | Q103 | S534 | G1082 |
| 85 | D104 | S535 | D1083 |
| 86 | L105 | L536 | L1084 |
| 87 | T106 | Y537 | K1085 |
| 88 | T107 | H538 | S1086 |
| 89 | Y108 | H539 | Y1087 |
| 90 | L109 | L540 | L1088 |
| 91 | D110 | H541 | R1089 |
| 92 | K111 | I542 | S1090 |
| 93 | V112 | I543 | L1091 |
| 94 | P115 | T545 | P1102 |
| 95 | G116 | K546 | P1103 |
| 96 | V117 | F547 | P1104 |
| 97 | P118 | E548 | T1105 |
| 98 | T119 | M549 | L1106 |
| 99 | E120 | I550 | Q1107 |
| 100 | T121 | K551 | E1108 |
| 101 | I122 | L552 | M1109 |
| 102 | K123 | I553 | I1110 |
| 103 | D124 | D554 | Q1111 |
| 104 | M125 | I555 | M1112 |
| 105 | M126 | A556 | A1113 |
| 106 | F127 | R557 | A1114 |

| Alignment | CDK6 (1G3N:A) | BRAF (4H58:C) | INSR (3ETA:A) |
|---|---|---|---|
| 107 | Q128 | Q558 | E1115 |
| 108 | L129 | T559 | I1116 |
| 109 | L130 | A560 | A1117 |
| 110 | R131 | Q561 | D1118 |
| 111 | G132 | G562 | G1119 |
| 112 | L133 | M563 | M1120 |
| 113 | D134 | D564 | A1121 |
| 114 | F135 | Y565 | Y1122 |
| 115 | L136 | L566 | L1123 |
| 116 | H137 | H567 | N1124 |
| 117 | S138 | A568 | A1125 |
| 118 | H139 | K569 | K1126 |
| 119 | R140 | S570 | K1127 |
| 120 | V141 | I571 | F1128 |
| 121 | V142 | I572 | V1129 |
| 122 | H143 | H573 | H1130 |
| 123 | R144 | R574 | R1131 |
| 124 | D145 | D575 | D1132 |
| 125 | L146 | L576 | L1133 |
| 126 | K147 | K577 | A1134 |
| 127 | P148 | S578 | A1135 |
| 128 | Q149 | N579 | R1136 |
| 129 | N150 | N580 | N1137 |
| 130 | I151 | I581 | C1138 |
| 131 | L152 | F582 | M1139 |
| 132 | V153 | L583 | V1140 |
| 133 | T154 | H584 | A1141 |
| 134 | S155 | E585 | H1142 |
| 135 | S156 | D586 | D1143 |
| 136 | G157 | L587 | F1144 |
| 137 | Q158 | T588 | T1145 |
| 138 | I159 | V589 | V1146 |
| 139 | K160 | K590 | K1147 |
| 140 | L161 | I591 | I1148 |
| 141 | A162 | G592 | G1149 |
| 142 | D163 | D593 | D1150 |
| 143 | F164 | F594 | F1151 |
| 144 | G165 | G595 | G1152 |
| 145 | L166 | L596 | M1153 |
| 146 | A167 | A597 | T1154 |
| 147 | R168 | T598 | R1155 |
| 148 | I169 | V599 | D1156 |

| Alignment | CDK6 (1G3N:A) | BRAF (4H58:C) | INSR (3ETA:A) |
|---|---|---|---|
| 149 | Y170 | K600 | I1157 |
| 150 | S171 | S601 | Y1158 |
| 151 | F172 | R602 | E1159 |
| 152 | Q173 | W603 | T1160 |
| 153 | M174 | S604 | D1161 |
| 154 | A175 | G605 | Y1162 |
| 155 | L176 | S606 | Y1163 |
| 156 | - | H607 | R1164 |
| 157 | T177 | E610 | G1167 |
| 158 | S178 | Q611 | K1168 |
| 159 | V179 | L612 | G1169 |
| 160 | V180 | S613 | L1170 |
| 161 | V181 | G614 | L1171 |
| 162 | T182 | S615 | P1172 |
| 163 | L183 | I616 | V1173 |
| 164 | W184 | L617 | R1174 |
| 165 | Y185 | W618 | W1175 |
| 166 | R186 | M619 | M1176 |
| 167 | A187 | A620 | A1177 |
| 168 | P188 | P621 | P1178 |
| 169 | E189 | E622 | E1179 |
| 170 | V190 | V623 | S1180 |
| 171 | L191 | I624 | L1181 |
| 172 | L192 | R625 | K1182 |
| 173 | Q193 | M626 | D1183 |
| 174 | S194 | N630 | G1184 |
| 175 | S195 | P631 | V1185 |
| 176 | Y196 | Y632 | F1186 |
| 177 | A197 | S633 | T1187 |
| 178 | T198 | F634 | T1188 |
| 179 | P199 | Q635 | S1189 |
| 180 | V200 | S636 | S1190 |
| 181 | D201 | D637 | D1191 |
| 182 | L202 | V638 | M1192 |
| 183 | W203 | Y639 | W1193 |
| 184 | S204 | A640 | S1194 |
| 185 | V205 | F641 | F1195 |
| 186 | G206 | G642 | G1196 |
| 187 | C207 | I643 | V1197 |
| 188 | I208 | V644 | V1198 |
| 189 | F209 | L645 | L1199 |
| 190 | A210 | Y646 | W1200 |

| Alignment | CDK6 (1G3N:A) | BRAF (4H58:C) | INSR (3ETA:A) |
|---|---|---|---|
| 191 | E211 | E647 | E1201 |
| 192 | M212 | L648 | I1202 |
| 193 | F213 | M649 | T1203 |
| 194 | R214 | T650 | S1204 |
| 195 | R215 | G651 | L1205 |
| 196 | K216 | Q652 | E1207 |
| 197 | P217 | L653 | Q1208 |
| 198 | L218 | P654 | P1209 |
| 199 | F219 | Y655 | Y1210 |
| 200 | R220 | S656 | Q1211 |
| 201 | G221 | N657 | G1212 |
| 202 | S222 | N659 | L1213 |
| 203 | S223 | N660 | S1214 |
| 204 | D224 | R661 | N1215 |
| 205 | V225 | D662 | E1216 |
| 206 | D226 | Q663 | Q1217 |
| 207 | Q227 | I664 | V1218 |
| 208 | L228 | I665 | L1219 |
| 209 | G229 | F666 | K1220 |
| 210 | K230 | M667 | F1221 |
| 211 | I231 | V668 | V1222 |
| 212 | L232 | G669 | M1223 |
| 213 | D233 | R670 | D1224 |
| 214 | V234 | G671 | G1225 |
| 215 | I235 | Y672 | G1226 |
| 216 | G236 | L673 | Y1227 |
| 217 | L237 | S674 | L1228 |
| 218 | P238 | P675 | D1229 |
| 219 | G239 | D676 | Q1230 |
| 220 | E240 | L677 | P1231 |
| 221 | V266 | R681 | - |
| 222 | T267 | S682 | D1232 |
| 223 | D268 | N683 | N1233 |
| 224 | I269 | C684 | C1234 |
| 225 | D270 | P685 | P1235 |
| 226 | E271 | K686 | E1236 |
| 227 | L272 | A687 | R1237 |
| 228 | G273 | M688 | V1238 |
| 229 | K274 | K689 | T1239 |
| 230 | D275 | R690 | D1240 |
| 231 | L276 | L691 | L1241 |
| 232 | L277 | M692 | M1242 |

| Alignment | CDK6 (1G3N:A) | BRAF (4H58:C) | INSR (3ETA:A) |
|---|---|---|---|
| 233 | L278 | A693 | R1243 |
| 234 | K279 | E694 | M1244 |
| 235 | C280 | C695 | C1245 |
| 236 | L281 | L696 | W1246 |
| 237 | T282 | K697 | Q1247 |
| 238 | F283 | K698 | F1248 |
| 239 | N284 | K699 | N1249 |
| 240 | P285 | R700 | P1250 |
| 241 | A286 | D701 | K1251 |
| 242 | K287 | E702 | M1252 |
| 243 | R288 | R703 | R1253 |
| 244 | I289 | P704 | P1254 |
| 245 | S290 | L705 | T1255 |
| 246 | A291 | F706 | F1256 |
| 247 | Y292 | P707 | L1257 |
| 248 | S293 | Q708 | E1258 |
| 249 | A294 | I709 | I1259 |
| 250 | L295 | L710 | V1260 |
| 251 | S296 | A711 | N1261 |
| 252 | H297 | S712 | L1262 |
| 253 | P298 | I713 | L1263 |
| 254 | Y299 | E714 | K1264 |
| 255 | F300 | - | - |
| 256 | Q301 | - | - |
| 257 | D302 | - | - |
| 258 | - | - | - |
| 259 | - | - | - |

**Table S3.** (*continued on next* page) 54 mutations totaling 108 FEP-calculated (alchemical) $\Delta G$s in the active $A$ and inactive basin $B$ calculated from the BAR estimator, along with the corresponding errors which are default output from FEP+. Each row represents a single pair alchemical transformation in basins $\mathbb{A}$ and $\mathbb{B}$. $\Delta\Delta G$ is calculated from the difference $\Delta G(\mathbb{B}) - \Delta G(\mathbb{A})$, where positive differences indicate a shift in stability towards the active basin, and negative values indicate a shift towards the inactive basin. These mutations were identified from Potts $\Delta\Delta T$ scans, where a total of 18 double mutations located far in the left and right-hand tails of the histograms in Figure 3A were broken down into two single mutants each. The corresponding values of $\Delta\Delta G_{reorg}$ plotted in Figure 3C were calculated by averaging over three different alchemical paths connecting the double mutant with the constituent single mutants, resulting in 36 additional "intermediate" mutations to close the thermodynamic cycle and calculate an estimate of the error in $\Delta\Delta G_{reorg}$ due to hysteresis. The location of single-point mutations in the active or inactive-state tertiary structure are annotated in the "Structural Motifs" column.

|  | Mutation | ΔG(A) | BAR error | ΔG(B) | BAR error | ΔΔG | BAR error | ΔΔG (Avg) | Hysteresis error | ΔΔT Potts | Structural Motif |
|---|---|---|---|---|---|---|---|---|---|---|---|
| INSR | M53Y/R123K | 7.98 | ±0.18 | 2.4 | ±0.14 | -5.58 | ±0.23 | -5.85 | ±0.58 | -1.9 | - |
|  | M53Y | -0.13 | ±0.14 | -3.76 | ±0.13 | -3.63 | ±0.19 | -3.88 | ±0.58 | -0.94 | R-spine |
|  | R123K | 9.08 | ±0.11 | 6.45 | ±0.07 | -2.63 | ±0.13 | -2.21 | ±0.58 | -0.99 | RD-pocket |
|  | L115A/F143L | 1.8 | ±0.12 | -2.75 | ±0.13 | -4.55 | ±0.18 | -4.19 | ±0.43 | -2.6 | - |
|  | L115A | -2.96 | ±0.08 | -4.42 | ±0.19 | -1.46 | ±0.21 | -1.86 | ±0.43 | -0.02 | R-spine |
|  | F143L | 4.5 | ±0.10 | 1.63 | ±0.13 | -2.87 | ±0.16 | -2.9 | ±0.43 | -0.49 | R-spine |
|  | F120I/F143Y | -1.37 | ±0.10 | -5.11 | ±0.16 | -3.74 | ±0.19 | -3.36 | ±0.35 | -2.1 | - |
|  | F120I | 5.96 | ±0.08 | 5.84 | ±0.12 | -0.12 | ±0.14 | -0.29 | ±0.35 | -1.01 | N-term. anchor |
|  | F143Y | -7.89 | ±0.08 | -11.04 | ±0.12 | -3.15 | ±0.14 | -3.31 | ±0.35 | -1 | R-spine |
|  | M145F/Y154L | 20.07 | ±0.10 | 23.49 | ±0.17 | 3.42 | ±0.20 | 2.93 | ±0.45 | 1.9 | - |
|  | M145F | 7.58 | ±0.10 | 8.16 | ±0.11 | 0.58 | ±0.15 | 0.67 | ±0.45 | 0.74 | N-term. anchor |
|  | Y154L | 12.31 | ±0.08 | 15.26 | ±0.12 | 2.95 | ±0.14 | 3.26 | ±0.45 | 1.11 | Substrate mimic |
|  | R123L/Y154F | 16.07 | ±0.14 | 18.81 | ±0.13 | 2.74 | ±0.19 | 2.85 | ±0.15 | 2.1 | - |
|  | R123L | 4.94 | ±0.11 | 5.2 | ±0.08 | 0.26 | ±0.14 | 0.27 | ±0.15 | 0.42 | RD-pocket |
|  | Y154F | 10.31 | ±0.11 | 13.12 | ±0.22 | 2.81 | ±0.25 | 2.65 | ±0.15 | 1.02 | Substrate mimic |
|  | R147A/Y154V | 3.61 | ±0.07 | 11.28 | ±0.19 | 7.67 | ±0.20 | 6.57 | ±1.07 | 1.4 | - |
|  | R147A | -7.1 | ±0.05 | -2.78 | ±0.19 | 4.32 | ±0.20 | 4.95 | ±1.07 | 0.36 | RD-pocket |
|  | Y154V | 10.81 | ±0.07 | 13.82 | ±0.22 | 3.01 | ±0.23 | 3.58 | ±1.07 | 1.05 | Substrate mimic |

| | Mutation | ΔG(A) | BAR error | ΔG(B) | BAR error | ΔΔG | BAR error | ΔΔG (Avg) | Hysteresis error | ΔΔT Potts | Structural Motif |
|---|---|---|---|---|---|---|---|---|---|---|---|
| BRAF | V52L/F143L | 3.68 | ±0.10 | 4.5 | ±0.10 | 0.82 | ±0.14 | 1.19 | ±0.40 | 2.3 | - |
| | V52L | 0.8 | ±0.11 | -0.16 | ±0.08 | -0.96 | ±0.14 | -0.97 | ±0.40 | 0.8 | N-term. anchor |
| | F143L | 3.77 | ±0.07 | 5.01 | ±0.11 | 1.24 | ±0.13 | 0.89 | ±0.40 | 0.33 | R-spine |
| | F131V/F178E | -150.16 | ±0.11 | -146.39 | ±0.09 | 3.77 | ±0.14 | 3.83 | ±0.55 | 2.4 | - |
| | F131V | 3.23 | ±0.09 | 6.56 | ±0.08 | 3.33 | ±0.12 | 2.83 | ±0.55 | 1.24 | - |
| | F178E | -153.79 | ±0.07 | -152.9 | ±0.06 | 0.89 | ±0.09 | 1.26 | ±0.55 | 1.11 | αEF-αF loop |
| | F131A/E157T | 23.79 | ±0.15 | 24.5 | ±0.12 | 0.71 | ±0.19 | 0.65 | ±0.06 | 2.7 | - |
| | F131A | -2.35 | ±0.07 | -1.2 | ±0.08 | 1.15 | ±0.11 | 1.18 | ±0.06 | 1.18 | - |
| | E157T | 26.33 | ±0.13 | 25.9 | ±0.10 | -0.43 | ±0.16 | -0.41 | ±0.06 | 1.53 | - |
| | I120Y/L145Y | -29.56 | ±0.10 | -28.89 | ±0.13 | 0.67 | ±0.16 | 0.77 | ±0.47 | -2.7 | - |
| | I120Y | -16.15 | ±0.09 | -16.65 | ±0.10 | -0.5 | ±0.13 | -0.18 | ±0.47 | -1.54 | N-term. anchor |
| | L145Y | -13.53 | ±0.11 | -12.4 | ±0.09 | 1.13 | ±0.14 | 0.71 | ±0.47 | -1.19 | N-term. anchor |
| | L53Y/I120R | -146.69 | ±0.14 | -151.44 | ±0.15 | -4.75 | ±0.21 | -4.14 | ±0.53 | -3.1 | - |
| | L53Y | -12.44 | ±0.08 | -11.98 | ±0.09 | 0.46 | ±0.12 | 0.35 | ±0.53 | -1.34 | R-spine |
| | I120R | -134.66 | ±0.09 | -138.93 | ±0.09 | -4.27 | ±0.13 | -4.6 | ±0.53 | -1.71 | N-term. anchor |
| | L53F/A146T | 11.99 | ±0.15 | 8.59 | ±0.16 | -3.4 | ±0.22 | -2.96 | ±0.90 | -2.9 | - |
| | L53F | -0.33 | ±0.09 | -0.66 | ±0.12 | -0.33 | ±0.15 | 0.18 | ±0.90 | -1.17 | R-spine |
| | A146T | 12.28 | ±0.10 | 9.61 | ±0.08 | -2.67 | ±0.13 | -3.41 | ±0.90 | -2.88 | N-term. anchor |

| | Mutation | ΔG(A) | BAR error | ΔG(B) | BAR error | ΔΔG | BAR error | ΔΔG (Avg) | Hysteresis error | ΔΔT Potts | Structural Motif |
|---|---|---|---|---|---|---|---|---|---|---|---|
| CDK6 | Y14L/T162Y | 1.3 | ±0.13 | 8.04 | ±0.19 | 6.74 | ±0.23 | 6.5 | ±0.76 | 3.1 | - |
| | Y14L | 13.87 | ±0.07 | 15.18 | ±0.12 | 1.31 | ±0.13 | 1.9 | ±0.76 | 1.63 | G-loop |
| | T162Y | -12.9 | ±0.11 | -7.81 | ±0.14 | 5.09 | ±0.18 | 4.59 | ±0.76 | 1.55 | C-term. anchor |
| | Y149L/V159F | 11.16 | ±0.11 | 14.49 | ±0.12 | 3.33 | ±0.16 | 4.14 | ±0.89 | 3.1 | - |
| | Y149L | 11.78 | ±0.08 | 15.32 | ±0.06 | 3.54 | ±0.10 | 3.64 | ±0.89 | 2.29 | N-term. anchor |
| | V159F | -0.74 | ±0.07 | 1.19 | ±0.08 | 1.93 | ±0.11 | 1.2 | ±0.89 | 0.67 | RD-pocket |
| | R147N/L163K | -39.64 | ±0.16 | -34.6 | ±0.10 | 5.05 | ±0.19 | 4.83 | ±0.38 | 2.5 | - |
| | R147N | -42.48 | ±0.11 | -41.08 | ±0.06 | 1.41 | ±0.12 | 1.18 | ±0.38 | 1.31 | RD-pocket |
| | L163K | -130.41 | ±0.08 | -127.15 | ±0.06 | 3.27 | ±0.10 | 3.59 | ±0.38 | 1.15 | C-term. anchor |
| | V120L/V160F | 2.76 | ±0.12 | -1.37 | ±0.06 | -4.13 | ±0.13 | -4.84 | ±0.75 | -5.2 | - |
| | V120L | 2.63 | ±0.08 | -0.51 | ±0.05 | -3.14 | ±0.10 | -3.08 | ±0.75 | -2.46 | N-term. anchor |
| | V160F | 0.57 | ±0.12 | -1.13 | ±0.06 | -1.7 | ±0.14 | -1.02 | ±0.75 | -2.77 | RD-pocket |
| | I148D/V160R | -153.7 | ±0.19 | -161.8 | ±0.05 | -8.1 | ±0.20 | -8.71 | ±3.07 | -5.1 | - |
| | I148D | -157.59 | ±0.06 | -159.99 | ±0.02 | -2.4 | ±0.06 | -1.31 | ±3.07 | -2.07 | N-term. anchor |
| | V160R | -131.94 | ±0.34 | -134.58 | ±0.04 | -2.64 | ±0.34 | -5.21 | ±3.07 | -3.11 | C-term. anchor |
| | R123K/Y176I | 27.85 | ±0.17 | 23.65 | ±0.16 | -4.2 | ±0.23 | -4.29 | ±0.41 | -6.3 | - |
| | R123K | 9.99 | ±0.12 | 5.44 | ±0.06 | -4.55 | ±0.14 | -4.24 | ±0.41 | -4.09 | RD-pocket |
| | Y176I | 17.81 | ±0.17 | 17.29 | ±0.13 | -0.52 | ±0.21 | -0.84 | ±0.41 | -2.3 | αEF-αF loop |
